# A spatiotemporal atlas of orchiectomy-induced androgen deprivation-mediated modulation of cellular composition and gene expression in the mouse prostate

**DOI:** 10.1101/2025.07.11.664414

**Authors:** Greg Shelley, Allison May, Tyler Robinson, Jinlu Dai, Sethu Pitchiaya, Evan T. Keller

## Abstract

Androgen deprivation therapy (ADT) remains a cornerstone in the treatment of prostate cancer (PCa), yet most tumors eventually develop resistance. Murine models are widely used to study PCa progression and ADT response, but a detailed understanding of the prostate’s biological response to androgen deprivation in these models is lacking. Here, we present a comprehensive spatiotemporal analysis of cellular and transcriptional dynamics in the mouse prostate following orchiectomy (ORX)-induced androgen deprivation. We observed progressive involution across all prostate lobes (dorsal, ventral, lateral, and anterior) and distinct lobe-specific temporal gene expression changes post-ORX. Immune cell infiltration markedly increased over time, highlighting a shift in the prostate’s cellular landscape. Single-cell RNA sequencing uncovered a previously undescribed fibroblast subtype—termed ORX-induced fibroblast (OIF)— characterized by high expression of Wnt2, Rorb, and Wif1, with distinct spatial localization. Pathway analysis revealed upregulation of amide and peptide binding functions, alongside suppression of peptidase and endopeptidase activity. Furthermore, dynamic changes in ligand–receptor interactions across lobes underscored the evolving intercellular communication in the post-ORX prostate. By integrating spatial transcriptomics with single-cell profiling, our study generates a high-resolution atlas of the murine prostate’s response to androgen deprivation. These findings provide a foundational resource for interpreting ADT responses in preclinical models of PCa.

## Introduction

Prostate cancer is the second most common cause of cancer-related death of men in the United States. When initially identified, most prostate cancers are dependent on androgens for growth leading to the clinical use of androgen deprivation therapy (ADT) through surgical or chemical means. While many patients initially respond to ADT, the majority of patients develop resistance to ADT and develop what is termed castrate-resistance prostate cancer (CRPC). Thus, understanding the mechanisms through which men progress to CRPC is an opportunity to identify therapeutic candidates to overcome resistance to hormonal therapy and improve patient care.

Although mice rarely spontaneously develop prostate cancer, murine models have long been used to study prostate cancer [1–3] including both orthotopically-implanted xenografts of human prostate cancer cells [4] and genetically engineered mice (GEM). An important aspect of some of these models is the ability to recapitulate their dependence on androgen for growth and subsequently the development of CRPC. Accordingly, defining how the mouse prostate responds to androgen deprivation is important to understanding how this response could impact orthotopic prostate xenografts and primary tumors in GEM murine models of prostate cancer.

There are multiple studies of the mouse prostate following experimental decrease of androgen levels [5–7]. Following androgen deprivation, the mouse prostate involutes with the loss of internal organization and the loss of many of the histological features associated with each of the lobes [8–11]. Within the prostate, many of the genes are androgen dependent and their expression is downregulated after castration [6, 7, 12–14], though little is known of the function of many of these genes. For example, β-microseminoprotein (Msmb) has been widely reported as being androgen dependent and to be expressed in the mouse prostate, however, although it is identified as a member of the immunoglobulin binding factor family [15, 16], its function in the prostate is unknown. Similarly, the function of probasin, a gene that is expressed in the mouse prostate and is the source of the prostate specific Pb promoter, is a protein that belongs to the lipocalin superfamily, but it’s functional role in the prostate is also unknown [17]. These studies have primarily been performed on bulk prostate tissues and thus are limited in their ability to identify gene expression changes within specific cell subpopulations. Identifying the cell subpopulations in which gene expression changes has the potential to facilitate defining the genes functions.

A challenge of modeling human prostate cancer in the mouse is the difference between the anatomy of the mouse and human prostate [18]. Specifically, the human prostate consists of one lobe that is composed of the central, transition and peripheral zones [19–22]. In contrast, the mouse prostate consists of four lobes, the anterior, dorsal, lateral and ventral, which are spatially distinct, each having their own morphology and histological features, such as position of nuclei with the cell, cytoplasmic granularity, infoldings and size of the acini [23]. The dorsal and lateral lobes have often been combined for research purposes as the dorsolateral (DL) [21–26], though recent studies have demonstrated that these are significantly different in terms of gene expression [27, 28]. Limited work has been performed on gene expression within the human prostate and how the different zones relate [29, 30], but it has been suggested that the murine DL prostate is most similar to the peripheral human prostate

[23]. Due to these multiple lobes and structural differences within the mouse prostate, the ability to clearly define the androgen deprivation-induced changes in cellular composition and gene expression within each lobe-type and further the spatial relationships between the cells and changes in the spatially-defined cellular niches would provide enhanced information towards an overall understanding of androgen deprivation on the prostate.

To determine information regarding different prostate lobes, there have been single cell transcriptomic studies [27, 28, 31, 32]. However, these did not explore changes following androgen deprivation. Furthermore, they were performed on tissues that had been made into cell suspensions so that the spatial relationships between cells could not be well-defined. To provide spatial context to cellular and gene expression changes in response to androgen-deprivation in the mouse prostate we previously performed spatial transcriptomics of mouse prostates on day 0 and day 15 post-Orx and explored changes that could impact androgen deprivation therapy for prostate cancer [33]. In the current manuscript, we explored the temporal nature of the changes in further detail through incorporating day 10 and day 20 post-orchiectomy and characterizing the temporal changes in the individual prostate lobes.

## Materials and Methods

### Mice

All animal work was performed with approval and in accordance with the University of Michigan Institutional Animal Care and Use Committee (IACUC). Male C57BL/6 mice 10 to 18 weeks old were either purchased from The Jackson Laboratory (Stock No. 00064; Bar Harbor, ME), or bred in-house from purchased mice. Mice were sham orchiectomized (incision and closure) (Sham) or orchiectomized as described previously [34] and prostates harvested 10, 15 or 20 days later. (T10, T15, and T20, respectively).

Upon removal of seminal vesicle (SV) and adhering fats, dissected whole prostates were flash frozen in 10x10mm cryomolds (VWR) containing TissueTek OCT (VWR). Cryomolds were immersed over a steel beaker of isopentane (Sigma) in LN2 per Visium Spatial Transcriptomics protocols (10X Genomics, Pleasanton, CA). Cryomolds were stored sealed in -80C freezer until use. In some cases, dissected whole prostates were placed in histology cassettes (Fisher) and fixed in 10% Formaldehyde for 4 hours and paraffin embedded within 24 hrs. Tissues were either cryosectioned into 5 to 7µm thick sections for frozen tissues or 5um sections for paraffin embedded samples.

### Spatial transcriptomics

Visium spatial analysis was performed using the whole genome assay in quadruplet on mice FFPE samples from four timepoints for a total of sixteen sections on four slides as previously described for the Sham and T15 sections [33]. Tissue sections of each sample type were analyzed for RNA integrity prior to placement on Spatial Tissue Optimization slides. Optimization was performed only for samples with RIN ≥7. Spatial gene expression slides specific for either cryosections or FFPE sections were used (10X Genomics, Pleasanton, CA) were processed according to the manufacturer’s instructions (10x Genomics; CG000239 Visium Spatial Gene Expression User Guide Rev) or (CG000408 Visium Spatial Gene Expression for FFPE –Tissue Preparation Guide and CG000407 Visium Spatial Gene Expression for FFPE User Guide). Samples were stained lightly with H&E on the Spatial Gene Expression slides using a standard protocol (CG000160 Methanol Fixation, H&E Staining & Imaging for Visium Spatial Protocols) and (CG000409 Visium Spatial Gene Expression for FFPE – Deparaffinization, H&E Staining, Imaging & Decrosslinking) and images scanned at high resolution and then probed with Visium Spatial Gene Expression Slide & Reagents Kit for the cryofrozen samples and the Visium Mouse Transcriptome Probe Set (Visium Mouse Transcriptome Probe Set v1. 0) for the FFPE samples.

Prostates were harvested at Days 0, 10, 15 and 20 post-orchiectomy and subjected to Visium-based spatial transcriptomics or single cell transcriptomics (Supplemental Figure 1). For spatial transcriptomics four consecutive sections from each prostate were evaluated at each time point (Supplemental Figure 2a).

Quadruplicate sections were subjected to Visium spatial transcriptomics; however, there was variance in how much of the sectioned prostate was successfully placed on the Visium slide (Supplemental Figure 2a). This resulted in variance in the number of Visium spots for each replicate, although there was no significant difference in the number of genes detected per sample or per spot (Supplemental Figure 2b) among the samples from within each time point. Placement on the slide was an additional factor to consider when analyzing the results, and one section each for T15 and T20, was missing a group of spots on the UMAP that was seen in the other three replicates (Supplemental Figure 2c). These corresponded to areas of the prostate that were not within the Visium slide capture area (Supplemental Figure 2a first column T15 and T20). Overall, minor differences in placement had minimal effect on genes and clusters identified and the correlation coefficient between each section from the same mouse was ≥0.99 for our first three time points (Supplemental Figure 2d). Overall, other than section A on the T20 slide, the correlation coefficient was ≥0.90 between any two slides from the same day. Correlation coefficient between the different days; however, was significantly lower, demonstrating that the detected gene expression changed temporally (Supplemental Figure 2e).

### Preparation of single cell suspension

After harvest, prostates were washed and kept in cold, serum-free RPMI medium (GIBCO). A digestion solution was made from Multi Tissue Dissociation kit 1 (Miltenyi) for C tubes composed of 2. Thirty-five ml of serum-free RPMI, 100µl Enzyme D, 50µl Enzyme R, and 12.5 µl Enzyme A. The prostate was transferred to a low bind plate and 150µl of the digestion solution was added. The prostate tissue was cut into 2 to 4mm pieces within 30 sec. to 1 min and transferred to the C tube with the same scalpel. The remaining cell suspension was pipetted into the C tube. Dissociation was performed in a gentleMACS Octo-dissociator with heater on program 37°C_Multi_A for 41 minutes (Miltenyi). Post digestion, cells were filtered through a 70µm strainer placed over a 50ml tube, filter-rinsed and tube spun for 300g for 10 min at room temp. RBC Lysis was performed with 1 ml 1X RBC Lysis buffer (Miltenyi) within 2 min at room temperature, then diluted with cold RPMI + 0.04% BSA and spun for 6 min 300g at 4°C. The supernatant was removed completely. Cells were run through debris /dead cell removal kit (Miltenyi) to increase to over 80% viability. Final cell suspension was in PBS + 0.04% BSA, spun down for 6 min at 300g at. Trypan Blue staining was used to count viable cells and cell counts were obtained using a BioRad C100 and validated with a hemocytometer to confirm that debris and extracellular material was not being counted as dead cells.

### Sequencing

Libraries for scRNA-seq were prepared using the 10x Genomics Chromium Single Cell 3′ Library and Gel bead Kit v3.1 according to the manufacturer’s protocol (CG000315 ChromiumNextGEMSingleCell3 v3.1 Dual Index RevA) to prepare 4 libraries of 10000 cells each. They were sequenced with a requested sequencing depth of 20000 reads/cell. Visium sequencing depth varied depending on whether the whole genome assay (200m reads/Visium section) or probe set assay (100m reads/sample) was used. Sequencing was performed using the NovaSeq6000 (Illumina) with 300 cycles and paired end 150. Each sequenced library was demultiplexed to FASTQ files and aligned to reads to the mm10 mouse transcriptome, for the single cell with Cell Ranger 6. 1.2 and for the Spatial Space Ranger 1.2 and 1.3.

Spatial analysis. Initial processing.

Initial processing of both single cell and spatial data was performed with Seurat v4.3 [35, 36]. Samples were aggregated into a single object using SCTransform with Seurat reciprocal PCA method with 2000 anchors [37]. Slides were combined into a single Seurat object, either with all 16 slides, or with one slide from each timepoint selected as having the greatest proportion of the section on the Visium slide.

### Clustering

Dimensional reduction was performed by first identifying the top 2000 variable features and using these to run PCA. Neighbors and then clusters were identified using the standard Seurat workflow with resolution 0.5.

### Differential Analysis

Differential analysis was performed in Seurat using FindMarkers to compare between specified conditions and FindAllMarkers to identify general marker genes, using the default settings which applied a Wilcoxon Rank Sum test. Differentially expressed genes were filtered to exclude genes with less than 1.5-fold differential expression or qvalue of less than 0.05. For some analyses, differential expression was performed using LOUPE browser as this allowed quick selection of cells or spots that could then be compared against each other. While these differential values were not used for further evaluation.

### Cell type identification

Initially, cells were annotated manually by their histology. Histological features were used as the basis for clusters in LOUPE browser. The sham was annotated first, and the four prostate lobes, anterior, dorsal, lateral and ventral could be identified as well as the urethra, the muscle surrounding it and the SV. There were also a couple of regions that did not have the morphological features to be assigned to any of the other.

Differential analysis was performed on these, and the marker genes compared to previously identified markers for each of the prostate lobes. Clustering was also performed in Seurat with a resolution of 0.5 on each timepoint, which gave 7 clusters for each. This matched the number of clusters manually identified. To validate, the clusters identified in Seurat were overlayed on the histological identified cluster and differential analysis performed on the regions where the unsupervised clustering was different to the manual clustering to determine which had better identified the prostate lobes.

Annotation of the later time points was performed in a similar manner, though the marker genes for sham prostate often were only weakly expressed in later time points, so cross validation was additionally performed using marker genes for each lobe identified on each of the days.

### Deconvolution

As training data and to check the algorithm worked, we used CARD [38] which had the best performance of the methods we tested as it was able to predict the presence of more cell types in the spatial data. We used cell types as identified in the single cell data to deconvolute the spatial data for the Sham and Day 15.

### Single Cell Analysis. Initial Processing

For the single cell experiment, cells with fewer than 100 genes were excluded prior to processing. Cells were filtered based on consecutive filtering, firstly by removing any cell with greater than 20% mitochondria genes, then by removing cells with under 1000 counts. Cells were also removed with an upper threshold limit based on the distribution of the counts or filters, typically with cells that had over 3 times the mean excluded as possible doublets. Further tests for outliers were performed as percentage ribosomal, percentage of all transcripts comprised of the most expressed gene and genes per UMI, but few cells were outside the range. Additionally, Cell Ranger Aggr was used to combine the samples for visualization and manipulation in LOUPE Browser.

### Cell type identification

After clustering was performed in Seurat, differential expression was performed on the clusters and cell types designated according to the differentially expressed genes. For those cell groups that were not annotated this way, clustering was performed on these alone. For the prostate lobes, we used marker genes for each lobe that had been previously identified in healthy prostate [24, 27, 28].

The majority of cells that had not been identified initially were fibroblasts and these were analyzed according to markers for fibroblasts in healthy mouse prostate as identified in two previous papers [31, 32]. Finally, two clusters that could not be identified by the marker genes were given a designation according to the genes expressed in these cells but not expressed or expressed at very low levels in all other cells. This was sufficient to classify 20478 out of 20637 sham cells and 26187 out of 26873 orchiectomy cells. For the spatial, this method allowed for annotating all cell in the Sham, approximately 90% of the T10 and T15, but only 67% of the T20.

### Pathway analysis

Differentially expressed genes were used to determine upregulated gene pathways or terms. Gene ontology terms that were differentially regulated were calculated with the R package cluster profiler [39]. KEGG, Hallmark, Reactome and Wiki pathway terms were all identified, as well as cell type signature gene sets, obtained through the Msigdbr package [40, 41]. Further visualization was performed with the Pathview package [42].

### Ligand Receptor interactions

Ligand receptor interactions and signaling pathways were primarily identified using the CellChat R package [43].

### Statistical Analysis

For differential expression, p-values and fold change were calculated in Seurat using the default settings.

## Results

### Orchiectomy drives marked changes in structure, cell composition and gene expression in the mouse prostate

Orchiectomy is frequently used in mice to model androgen deprivation of human prostate cancer [3]. Accordingly, to examine the impact of androgen deprivation on the spatial biology of the prostate mice were orchiectomized (or sham orchiectomized as control). Prostates were then harvested on Days 0, 10, 15 and 20 post-orchiectomy (ie., Sham, T10, T15, and T20, respectively) and subjected to Visium-based spatial transcriptomics or single cell transcriptomics (Supplemental Figure 1). For spatial transcriptomics four consecutive sections from each prostate were evaluated at each time point (Supplemental Figure 2a, b).

We observed a decrease in the overall size of the prostate and SVs following orchiectomy (Figure 1a). A large component of the overall decrease was due to the loss of the SVs, which were mostly absent in T20 (Figure 1a, b). Histology revealed involution of all prostate gland structures as well as the SVs (Figure 1b). By T20, there was a marked loss of epithelial cells with the majority of remaining cells being stromal and a large component of connective tissue (Figure 1b).

**Figure 1.**
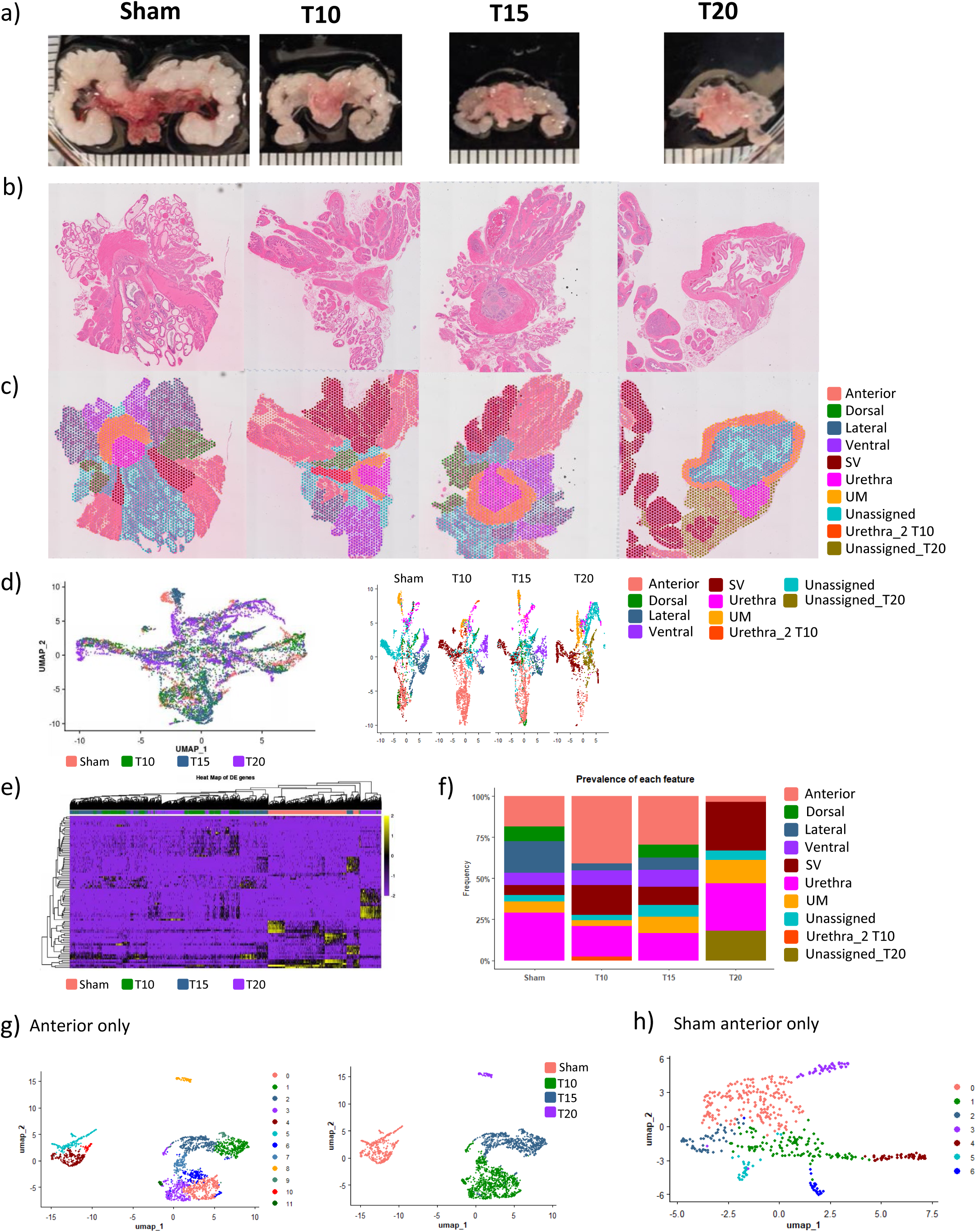
Spatial transcriptomics reveals cellular and transcriptomic spatially-defined orchiectomy-induced changes. Mice were subjected to sham surgery or orchiectomy (orx) and prostates collected and indicated timepoints (T in days). Prostates were then subjected to Visium spatial transcriptomics. (a) Whole prostates at each timepoint, scalebar = 1mm division. (b) Hematoxylin and eosin (H&E) stained prostate sections. External SVs were removed prior to embedding and samples stained. (c) Annotation of the prostate at each timepoint. Histology was used to broadly define the different regions, which was refined using clusters identified in Seurat. Spot assignation shown in legend. (d) Visualization of spots with Uniform Manifold Approximation and Projection (UMAP) of all samples combined (left UMAP) then split by timepoint and colored by annotation (right UMAP). (e) Heat map of the most differentially expressed genes at each timepoint. The top 20 upregulated genes and top 5 downregulated genes for each timepoint were chosen based on magnitude of fold change. Only genes detected in at least 25% of spots were included for the upregulated genes. (f) Stacked bar plot showing the prevalence of each feature at each time point. (g) The spots annotated as Anterior on each of the four timepoints were reanalyzed as a subset. UMAP of the anterior lobe only at each of the time points, colored by the cluster (left) or the timepoint (right). (h) UMAP of the anterior lobe in the Sham sample only at resolution 0.8.

To make use of the spatial nature of this data, we first identified the different features in the Sham prostate. We performed both manual clustering based on histology and unsupervised clustering based on gene expression using Seurat (Figure 1b, c, Supplemental Figure 2d). The final annotation was primarily based on histology, but in some instances, such as where to divide between the anterior and the SV, we were guided by clustering as performed in Seurat (Figure 1b, 1c). All prostate lobes were distinctively identified in addition to non-prostate structures such as the SV and urethra, the latter which was further divided into the inner urethra and the muscle surrounding it (Figure 1c). While the results were consistent between samples from the same time point, it was low between those of different timepoints (Supplemental Figure 2e, f).

Across the sections, we saw that number of counts and genes varied, both by timepoint and spatially (Supplemental Figure 3a, b), notably the muscle surrounding the urethra had a lower number of detected genes, but other features, such as one of the SVs in the Sham also had significantly lower counts (Supplemental Figure 3a).

Although there were regions on the spatial plot that had higher counts, these were fairly evenly distributed and did not represent any morphologically observable features.

We selected one slide from each timepoint that had the most complete representation of the tissue at that timepoint within the Visium slide area (Figure 1b) for the primary visual representation. We performed dimensional reduction on each of the samples individually and saw that the features appeared separately on the UMAP (Supplemental Figure 3c). In the Sham, the SV appeared as two separate groups of cells, which may have been a function of the lower observed UMI and gene counts, though they were part of the same cluster (Supplemental Figure 3c).

We validated these results by using previously identified marker genes from two single cell data sets (Crowley et al. and Graham et al. [27, 28]) and a bulk microarray dataset (Berquin et al. [24]) for the anterior, dorsal, and ventral lobes (Supplemental Figure 4). We found that while the genes identified by these previous groups in the single cell experiments for the lateral and ventral were both strongly expressed and specific to those regions in our data, the anterior and dorsal genes did not differentiate the anterior and dorsal as well. We also saw some of the Crowley anterior marker genes were primarily expressed in the urethra (Supplemental Figure 4). However, for each of the regions, we identified highly specific marker genes (Supplemental Table 1). Taken together with our ability to spatially map the genes and clusters, this increased our confidence that the marker genes accurately identify the prostate features.

Using the marker genes for each of the prostate lobes (Supplemental Table 1) we were then able to annotate the later timepoints. Based on a combination of morphology and gene expression, the anterior lobe was still present at T10 and T15, and at T20 (Figure 1b, c). A feature that expressed the genes identified by Berquin as dorsal (Supplemental figure 4) and expressed many SV-related genes, was present throughout the time course based on expression of these genes, though by T10 its appearance had changed significantly. By T20, none of the prostate lobes could be distinguished based on histology alone, and the urethra had significantly changed in appearance. Ultimately, by combining histology and gene expression, we were able to identify what remained of the different prostate lobes (Figure 1c, d). As with the Sham, features identified using histologically annotated versus unsupervised clustering in the T10 through T20 prostates largely overlapped indicating that the gene expression was able to robustly differentiate prostate lobes even after their involution.

### Orchiectomy induces varied cellular and gene expression changes among the different prostate lobes

After processing and integration of all sections, we performed dimensional reduction and observed a similar overall clustering pattern of the spots across the timepoints (Figure 1d). The spots for the different timepoints did not form discrete clusters but largely overlapped (Figure 1d, note the overall UMAP split out by time points), though there were groups of cells that were specific to each timepoint.

There were genes that were differentially expressed for each timepoint. A heatmap of these genes demonstrated that timepoints overall clustered separately, indicating that orchiectomy induced temporal changes in change expression (Figure 1e).

Post-orchiectomy, the prevalence of many features changed over time (Figure 1f). In the Sham, the anterior, dorsal and lateral lobes each represented approximately 20-22% of the prostate, while the ventral lobe represented approximately 10% of the prostate. The remainder of the slide was comprised of SV and urethra. In T10, the anterior composed a larger percentage of the slide (41%), and the lateral was reduced (Figure 1c, f) and by T20, neither the lateral or ventral were detected (Figure 1c, f). As some genes were lobe specific and the relative frequencies of the lobes changed, we would expect their prevalence to change over time, even if their expression was not downregulated at the cellular level.

As the prostate lobes changed temporally, we analyzed each lobe independently. The anterior lobe was present at all time points and through T15 and maintained sufficient morphological features to be identifiable from the H&E image (Figure 1c). We performed dimensional reduction of the anterior prostate alone and identified multiple clusters and then determined the temporal relationship of these clusters by labeling spots based on their timepoint (Figure 1g). We identified that each timepoint resolved as a unique cluster (Figure 1g) indicating that the cells composing the anterior prostate were transcriptionally different at each time point. For example, three of these clusters (Clusters 4, 5 and 10) were primarily in the anterior prostate of the sham, and conversely, this timepoint was composed primarily of these clusters (Figure 1g). Differentially upregulated genes for between these three clusters included: Cluster 4) Erich3, Ren1; Cluster 5) Pfkfb3, Crisp1; and Cluster 10) Upk3a, Pate7 (Supplemental Figure 5a).

To further refine composition of the anterior prostate we performed dimensional reduction on only the Sham timepoint of the anterior lobe and identified 7 clusters, (Figure 1h, Supplemental Figure 5b). Analysis of these markers showed that while there was spatial localization of the expression, they didn’t define spatial niches within the Anterior. However, there was some spatial localization, for example, cluster 2 genes showed higher dorsal expression and cluster 3 expressed in the unassigned region ventral to the urethra (Supplemental Figure 5c). This may be a consequence of the Visium technique in that each spot may contain 8-10 cells, so, especially round the interface between two regions, spots may contain multiple cell types. Some of these genes were also temporarily differentially expressed. For example, Retnla1 and C3, markers for cluster 1 were both expressed at higher levels in T10 and T15 and Ren1, a marker for cluster 0 (though it was seen in most anterior cells) was also widely expressed in T10 anterior, where it was also limited to the anterior prostate (Supplemental Figure 5c).

In the dorsal lobe (Supplemental figure 6a-c), each timepoint also grouped together on the UMAP, though unlike the anterior, the T15 was spatially distinct, rather than the Sham. There were subsets that had increased expression of 0) Dkki1, Kcgn2, 1) Map7d2, Tmef2 and 2) Kcnc1, Kcnip1. None of these mapped to particular areas within the dorsal lobe, though they did have spatial localization in other lobes, for example, Dkki1 was higher in the SV. The genes for cluster 1 were strongest in the SV-like tissue though they were also expressed at a higher level in the T10 dorsal. Taken together, these data show that the individual prostate lobes have orchiectomy-induced temporal-dependent transcriptional changes that are unique to each lobe.

### scRNAseq identifies cellularity differences during castration

To further investigate the cell types present in the prostate, we performed single cell RNA-seq on the prostates from Sham and T15 mice. Each specific prostate lobe epithelia clustered separately between Sham vs. T15 (Figure 2a). We used prostate lobe marker genes defined from the spatial analysis to assign the cells to the different prostate lobes, which made up 43.8% of cells in the Sham, and 16.3% of cells in the T15 mice (Figure 2b). This was due to marked reduction of all prostate lobes. The remainder of the cells were primarily fibroblasts (approximately 40% in both timepoints) and immune cells (10.7% of cells in sham and increased to 35.4% of cells in the immune), with a small proportion of other cells (Figure 2b-c). The single cell data mapped to the spatial data which demonstrates that the single cell data labeled the appropriate tissues.

**Figure 2.**
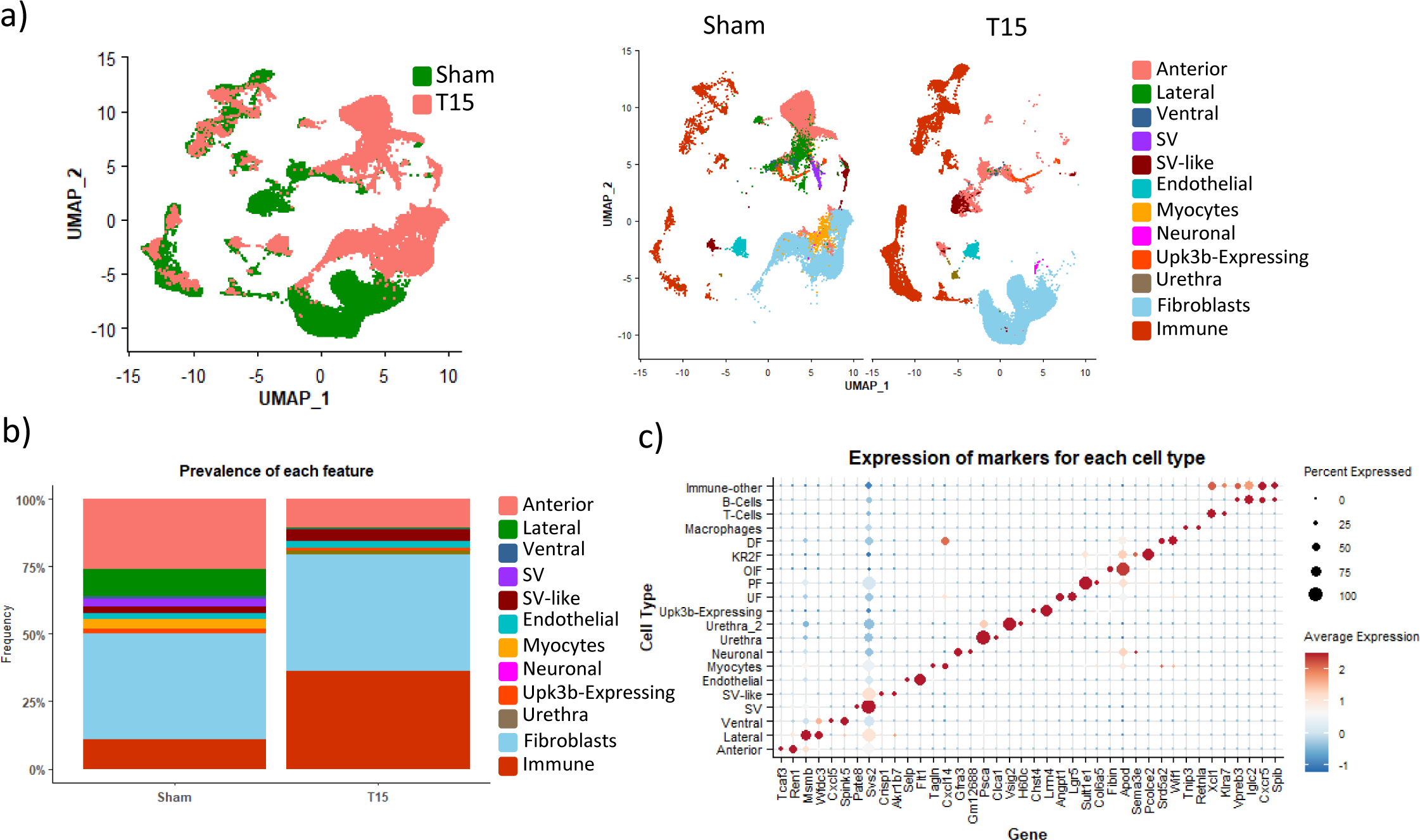
Single-cell RNA sequencing reveals cellular and transcriptome orchiectomy-induced changes. (a) UMAP of single cells from Sham or T15 mouse prostates: annotated by timepoint (left) then split by timepoint (Sham versus T15) and colored by cell annotation (right). (b) Stacked column of cell distribution in Sham vs T15. (c) Dot plot of cell type vs Marker genes. The top two differentially expressed genes for each identified cell type are shown

Similar to the prostate lobes, many of the fibroblasts clustered separately between Sham vs. T15. These were identified as different fibroblast subtypes based on previously published data [31, 32] (Figure 2c) indicating that both the different prostate lobe epithelia and the fibroblast populations underwent significant changes in gene expression post-orchiectomy. In contrast, the immune cells and endothelial cells from both timepoints clustered together, respectively (Figure 2a), suggesting that they were transcriptionally similar between time points.

### Single cell data reveals cell types not identified in the spatial data

In addition to the well-recognized cells in the prostate, the single cell analysis revealed rare cell types. Specifically, we identified small populations of myocytes, endothelial cells and neuronal cells (Figure 2b-c). An additional population of cells we identified we termed Upk3b-expressing as these cells expressed this gene, but it was only detected in 1.4% of the other cells. While this gene is urethral, other genes, such as Msln, Nkain4, C2 and Cavin2 were also detected at much higher percentage in these cells as opposed to the rest of the population, in both the Sham and Orx. Msln is associated with mesothelial cells lining the kidney and Cavin2 with endothelial cells. Thus, we were not confidently able to assign a specific cell type to the Upk3b-expressing cells. This population clustered separately on the UMAP and had little change between Sham and Orx.

### Orchiectomy is associated with an increased prevalence of immune cells in the prostate

Analysis of the scRNAseq data of Sham and T15 prostates revealed that the overall percentage of immune cells increased, from approximately 10% in the Sham to approximately 35% at T15 (Figure 3a). In the Sham, B-cells, T-cells and macrophages made up 1.1%, 3.3% and 6.4% of all cells respectively, whereas at T15, they made up 3%, 18.5% and 14.6% (Figure 3a). To better define changes in the immune cells, we analyzed just the immune cell subset in the scRNAseq data. We were also able to identify a number of immune cell subtypes based on expression of known markers, such as Cd8a for the Cd8 T-cells and Klrd1 and Nkg7 for NK-T-cells (Figure 3b). We also observed several subsets of macrophages, which we termed M1-like, M2-like and M2a-like, based on expression of markers associated with these macrophage subtypes, plus a fourth population that expressed macrophage markers, but not the ones specific to these subtypes, which we termed unknown macrophages (Figure 3b). We next performed dimensional reduction on the immune cell population itself for both time points (Figure 3c). This revealed a similar cluster pattern in the immune populations, albeit with different cell densities of Sham and T15 (Figure 3c). The frequency of the subtypes was altered between time points (Figure 3d). For example, cells identified as NK-T cells and CD8 T-cells represented an increased proportion of the immune population at T15 compared to the Sham. In contrast, the M1-like and M2-like macrophages represented a lower proportion at T15 compared to Sham; whereas, the proportions of M2a-like macrophages and other macrophage types did not differ across the time points (Figure 3d).

**Figure 3.**
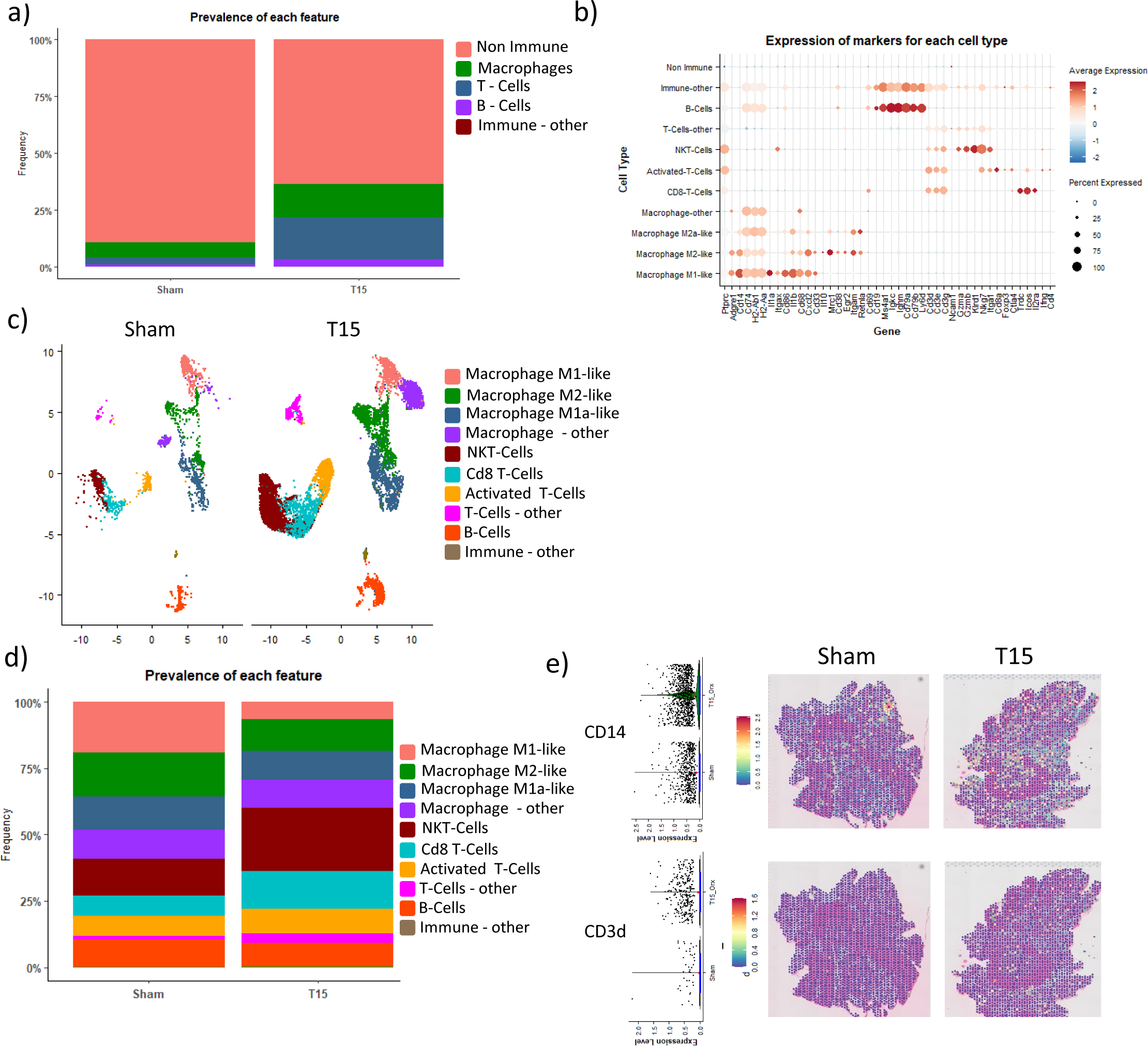
Androgens impact immune response. Prostates at time 0 (Sham) and 15 days post-orchiectomy (T15) were subjected to scRNAseq. The cells identified as immune cells in the scRNAseq analyses were selected and subjected to bioinformatic analysis. Subsets of T-cells, macrophages and B-cells were annotated based on the presence of markers associated with known immune cell subtypes. (a) Stacked column of immune or non-immune cell distribution by time point showing immune cell distribution in total single cell population prior to analyzing them as a subset. (b) Dot plot of markers for selected immune subtypes in the immune cells from the single cell data. (c) UMAP of the immune cells from the single cell data, split by the timepoint. (d) Stacked column of proportion of different immune cells identified as a percentage of the whole immune population T15. (e) Expression of Cd14 (top) and Cd3d (bottom) in the spatial data in Sham and T15.

Having determined that in the single cell data, both sham and post-orchiectomy prostate contained a large proportion of immune cells, we sought to identify these in the spatial data. We saw little expression of Cd3d in either Sham or T15, albeit it was higher in T15, and Cd14 was higher in the T15 (Figure 3e, Supplemental Figure 7a).

Interestingly, there was a concentrated focus of Cd14 expression within the lateral lobe of the Sham (Figure 3e). Other immune genes, such as Ptprc (general immune cell marker), Adgre1 (Macrophage marker), Cd3e (T-Cell marker), Cd19 and Cd79a (B-Cell markers) were detected as expected in the appropriate immune cells but only seen at low levels in the spatial data (Supplemental Figure 7b). An exception is Cd74, which was highly expressed in almost all spots (Supplemental Figure 7c). This was supported by the single cell data, in which it was seen in every cell type, and as many as 35% of the endothelial cells (Supplemental Figure 7a).

There are several published methods to estimate the likelihood that a spot contains different cell types, such as determining the module score in Seurat to using single cell expression data from other data sets to predict the identity of a spot, or to deconvolute spatial data using single cell data such as SPOTlight [44], CARD [38] and RCTD [45]. Other than an area of spots in the Sham that showed a high concentration of macrophages (Supplemental Figure 7d), in the same position as the spots with high Cd14 expression, we did not identify many spots predicted to contain a high percentage of immune cells using these methods.

### Orchiectomy promotes a novel fibroblast subtype

Although fibroblasts are known to be present in the kidney [31, 32], they were not identified as discrete spots in the spatial data which was likely due to them representing only a minor proportion of cells in each Visium spot (recall that each Visium spot is a combination of multiple cells). It has also previously been observed that fibroblast function is affected by AR levels in prostate cancer [46], which may have multiple subtypes [47]. In contrast to the spatial analysis, the single cell analysis did reveal fibroblasts (Figure 2a, b). Orchiectomy did not significantly alter the percentage of cells identified as fibroblasts, as fibroblasts represented 36% and 32% of all cells in the Sham and T15 prostates, respectively. We had previously seen that the fibroblasts in the Sham and the T15 appeared as largely discrete groups of cells on the UMAP comprised of several sub-clusters (Figure 2a) and this was maintained when we viewed them individually (Figure 4a).

**Figure 4.**
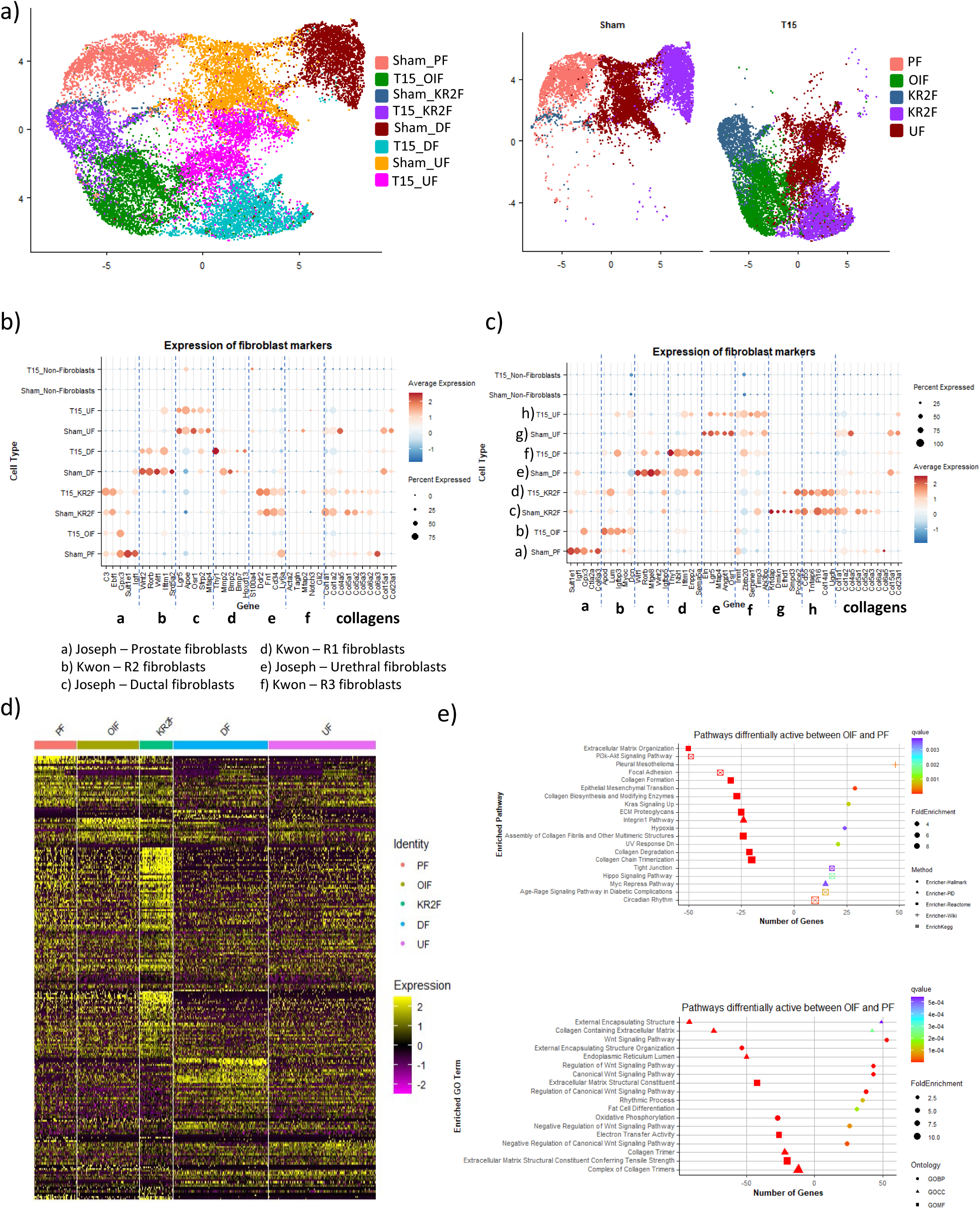
Orchiectomy impacts gene expression of fibroblast cells. Prostates at time 0 (Sham) and 15 days post-orchiectomy (T15) were subjected to scRNAseq. The cells identified as fibroblasts in the scRNAseq analyses were selected and subjected to bioinformatic analysis. These were annotated based on the expression of previously published fibroblast types seen in healthy murine prostate. (a) UMAP of fibroblasts from Sham and T15 combined (left) and UMAP of fibroblasts separated by time point (right). (b) Dot plot showing expression of marker genes for fibroblast types identified from previously published scRNAseq mouse prostates. Marker genes for the primary three fibroblast types identified as described in the text of each paper. (c) Dot plot showing expression of marker genes for each fibroblast type as identified in this paper. For each fibroblast type, the top 5 markers based on fold change were chosen in both Sham and Orx. (d) Heat map of the most differentially expressed genes in each of the fibroblast types PF, OIF, KR2F, DF and UF. For the heat map, the top 95 upregulated and 5 downregulated genes were chosen for each cluster, both as a single type (eg DF) or as how they separated between Sham and T15 (ie 8 cell types), for 203 genes. (e) ORA for top) pathways differentially active between OIF and PF, bottom) Go terms differentially regulated between OIF and PF.

The majority of previously published data on prostate fibroblasts has been in the context of prostate cancer [47–49], however, there are previous reports detailing different fibroblast subtypes in healthy prostate. Joseph at al. [31] identified three predominant fibroblast types, which they termed prostate fibroblasts, ductal fibroblast and urethral fibroblasts. Kwon et al. [32] also identified three primary types, which they termed R1, R2 and R3. Based on the marker gene expression identified by these authors in our data, the Kwon-R1 fibroblasts were equivalent to Joseph-Ductal fibroblasts and were called Ductal fibroblasts (DF) in our samples (Figure 4b). We saw common gene expression between the Kwon-R2 type fibroblasts and the Joseph-Prostate fibroblasts in our data, but there were sufficient differences, especially the expression of collagen related genes, to be confident that these were actually two different types of fibroblasts (Figure 4b,c), which we termed Joseph Prostate fibroblast (PF) and Kwon R2 fibroblasts (KR2F), though there were few of the KR2F in the Sham. For each of these types of fibroblasts, i.e. DF, Urethral (UF) and KR2F that were present in both the Sham and the T15, they were distinct on the UMAP and to varying extents, had a different gene expression profile and marker genes. This was least clear for the KR2F, which did have genes unique to the Sham, but the genes that were markers in the Orx were also present in the Sham, although this may be because of the limited numbers in Sham (Figure 4b).

We then determined the top 5 differential genes (based on magnitude of expression) for each fibroblast cell type and mapped the cells to data from the previous publications. This allowed us to identify differences in gene expression between Sham and T15 prostates for similar cell types. For example, Mfge8 was a marker for the DFs in the Sham, but not in the T15, while conversely, Thy1 was seen in the Sham DFs, but not the T15 (Figure 4c).

Based on the gene signature of the previously identified PFs, we did not observe the PFs in the T15 sample, but did see a novel subset, which we termed "Orchiectomy-induced fibroblasts" (OIF). Although some genes, such as Gpx3a were only expressed at high levels in these and the PFs, the overall gene expression patterns were different, and they did not express many of the PF marker genes (Figure 4b, 4c). Conversely, the top marker genes for these OIFs showed poor expression in the PFs. Overall, the top marker genes for each fibroblast cell type showed high specificity, and a heat map of these showed clearly distinct patterns across the different type (Figure 4d). The differential gene expression observed in these fibroblast subsets gives little clue as to their function, except the DFs are likely involved in Wnt signaling as Wnt2, Rorb and Wif1 are the most upregulated genes in these fibroblasts.

We also wanted to better understand the differences between the PFs and OIFs, so ran pathway analysis on these two cell types. We observed that gene ontology terms or pathways related to Wnt signaling were upregulated in the OIF, and while some collagen terms, such as "Collagen containing extracellular matrix" had genes upregulated in both, other collagen terms, such as relating to collagen trimers or tensile strength were more prevalent in the PFs (Figure 4e), suggesting that the collagen in the Sham was more highly organized. We also saw that genes regulated by MDM2, which is involved with P53 regulation [50] were enriched in the OIFs (Not shown).

### Orchiectomy induces changes of expression in multiple genes

Orchiectomy induced temporal downregulation of multiple genes in the prostate, both as overall expression (fold change) (Figure 5a, Supplemental Table 2) and as a change in the percentage of spots expressing a gene, or as a combination of the two. While we compared lobe-level changes for all time points, we focused on the T15 vs Sham differential expression markers as we had generated both single cell and spatial data for these time points. For downregulated genes, Spink1 which had the largest downregulation, also had a decrease in the percentage of spots expressing a gene (Figure 5b, c), from 100% in Sham to just 18% at T15. Other genes, such as Pbsn and Svs5 were detected in almost all cells at both time-points but still had a large decrease in average expression (Supplemental Table 2).

**Figure 5.**
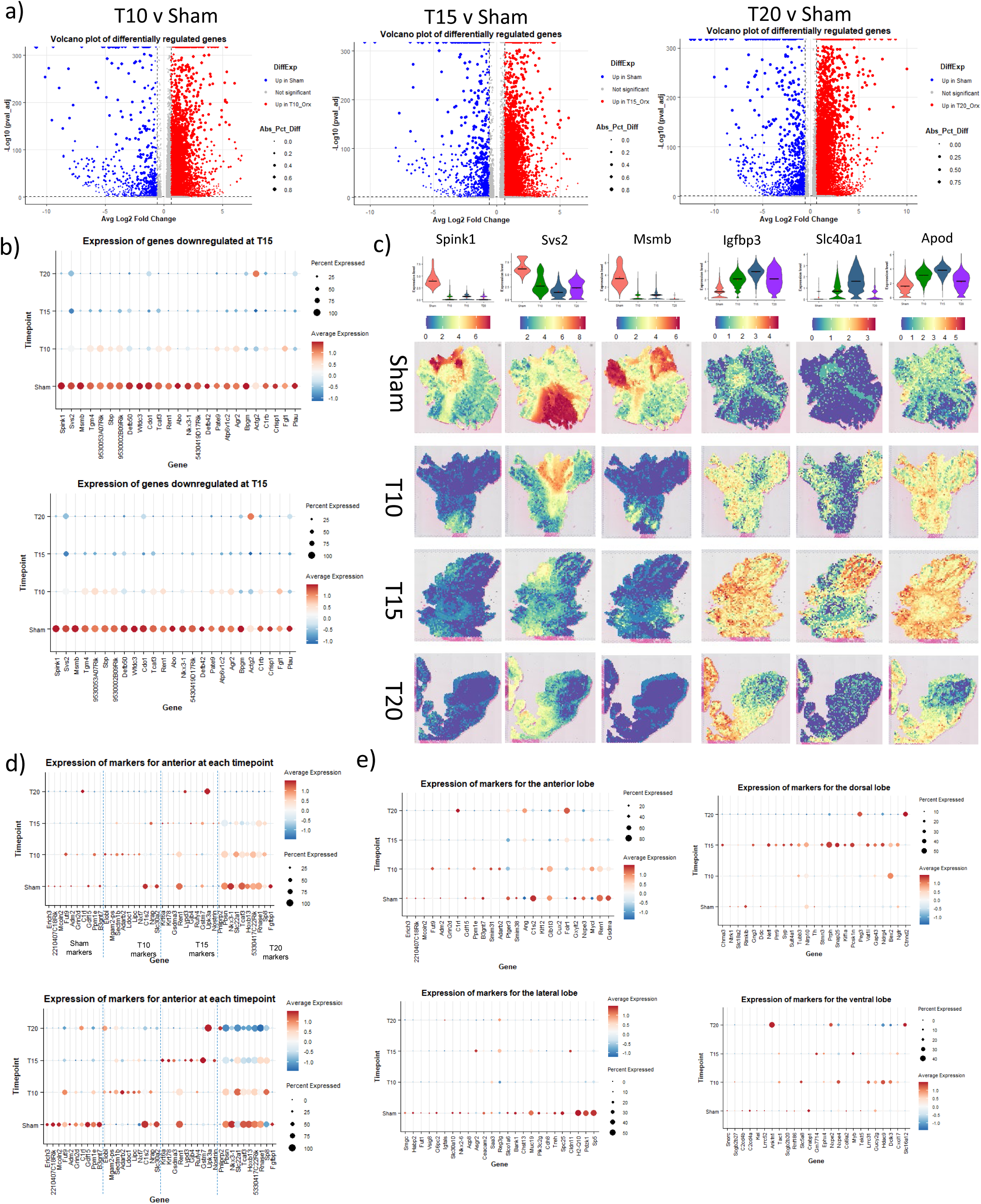
Changes in gene expression. (a) Volcano plot of differentially expressed genes for each of the timepoints compared to gene expression in the sham prostate. Genes with adjusted p-value <0.05 and absolute log2 fold change >0.6 colored, downregulated in blue, upregulated in red. Difference in expression indicated by size of dot. (b) Dot plot of expression of genes downregulated (upper plot) and upregulated (lower plot) at T15 compared to Sham. (c) Expression of selected highly differentially expressed genes between T15 and Sham. A violin plot shows the overall distribution of the gene’s expression over time (upper) and the images (lower) show the spatial distribution of the gene’s expression for each time point in the prostate. The first 3 genes are examples of genes downregulated at T15; whereas, the last 3 genes are examples of upregulated genes. (d) Dot plot of expression markers identified as the 10 most upregulated in the anterior region at each of the four timepoints. Shown are all spots (upper plot) and Anterior only (lower plot). (e) Dot plot showing expression of marker genes for each lobe as identified in the Sham prostate, at different timepoints.

The top 20 genes differentially downregulated in T15 compared to the sham were also reduced in expression or frequency in T10 compared to sham, and mostly further reduced in T20, albeit some genes, such as Actg2, were increased at T20 compared to T15 (Figure 5b). The pattern for the upregulated genes was similar, though somewhat more variable. Most were absent or present at low levels in the sham, and increased expression at T10 with continuing increases through either T15 or T20 (Figure 5b, 5c, Supplemental Figure 8). The gene downregulation may be simply due to genes no longer being activated in the absence of androgens, whereas the upregulated genes may be due in part to activation secondary to androgen loss or in part due to new pathways that are activated or the presence of additional cell types within the tissues, such as the recruitment and increase of immune cells that could result in the presence of immune related genes. Change in frequency of the different cell types over time accounts for some of the gene expression changes, but for the downregulated genes, their expression level within each cell type was decreased (Figure 5c) demonstrating specific downregulation of the gene expression itself.

### Spatial localization of gene expression

We next wanted to determine if genes that were temporally differentially expressed were spatially localized. Of the highly downregulated genes, Spink1 and Msmb were almost entirely downregulated; whereas, Svs2 expression was partially maintained in the SV (Figure 5c). Other genes, such as Svs5 (SV secretory protein 5) showed a similar pattern but there were genes such as Pate14 (prostate and testis expressed 14) that were initially expressed in the dorsal but increased in the SV (Supplemental Figure 8b). The genes that were upregulated also tended to show a pattern of preferential spatial expression. Apod (Apolipoprotein D) and Igfbp3 (Insulin-like growth factor-binding protein 3) were at lower levels in the Sham, preferentially seen in the anterior in T10 and throughout the prostate in T15 (Figure 5c), further suggesting that the different lobes are altered differently. Other genes were not expressed in the Sham, but were expressed at later time points, for example Pate7, was seen in the SV from T10 onwards (Supplemental Figure 8a).

Another gene that had temporally increased expression is clusterin (Clu), which encodes for a heat shock protein known to activate a cell survival program [51] and has been identified as protecting cells from androgen depletion [52, 53]. It is present in low levels in the Sham and observed at higher levels in a small group of spots that also have increased levels of many immune genes and so likely contain immune cells (Supplemental Figure 8a). In T10 through T20, its highest expression is in the SV. However, these spots do not show higher level of genes known to interact with clusterin or to be involved in apoptosis inhibition, so the role that clusterin plays in these cells is unclear (data not shown).

Finally, some genes show a mixed pattern of expression, for example, Nupr1 is a ventral gene in the Sham and T10, but in T15, and to a lesser degree T20, it is expressed in the Anterior. 9530002B09Rik is expressed throughout the 4 prostate lobes in the Sham, but by T10 and T15, it is predominantly seen in the Anterior, though some expression in the other lobes is maintained (Supplemental Figure 8a). The genes that did not show a consistent pattern over the four time points were generally spatially localized within the same region; for example, Actg2 was highly expressed in the urethra muscle in T20 but was much lower in the muscle on the earlier time points (Supplemental Figure 8a). Other actins had a similar pattern.

Taken together, these data demonstrate that orchiectemy induces different gene patterns in different prostate lobes and it is critical to look at each gene temporally and spatially in order to delineate its expression pattern.

### Mapping single cell data to Visium data provides additional spatial gene expression information

Having identified cell types in the single cell analysis that we had not been able to detect in the original analysis of the Visium spatial data, we attempted to see if we were able to map these cells to the spatial data based on their expression profiles. We used the single cells as training data and deconvoluted using CARD [38]. We included all cell types and found that for some, such as the anterior, the match was very good, with the spots in the region we had annotated as being anterior predicted to contain typically 80-100% anterior cells (Supplemental Figure 9). The other lobes did not have quite as high a match, but spatially, the spots predicted to contain that type of cell did match the expected area. The cellular annotation for the T15 showed a similar pattern

Having verified that the marker genes from the single cell were useful to annotate the spatial data, we also evaluated for the other cell types. One, that we had termed "myocyte" due to the presence of several differentially upregulated smooth muscle and myosin genes, was most strongly seen in the urethra, though a low percentage of the SV-like spots also contained cells expressing these genes (Supplemental Figure 9).

In the T15 however, a small proportion of the spots designated as ventral were predicted to be both partially anterior and partially ventral. Other features that mapped spatially in the T15 were the OIFs and the unknown macrophage type cells, both of which seemed to be present in the urethra muscle region. The OIFs were also present in the tissue surrounding the urethra (Supplemental Figure 9).

### Orchiectomy induces lobe-specific changes in gene expression

As much of the differential expression could be explained by changes in prevalence of different features or lobes, we looked at differential expression within each lobe. We found that the marker genes changed over time (Figure 5d). For example, the genes that most strongly marked the anterior in the Sham were expressed much lower in T10, and many were expressed at very low levels in a much smaller proportion of cells. Similarly, the genes that most strongly differentiated the anterior at T15, were generally poor markers for anterior in the Sham, and many were not even differentially expressed at that time. Unlike the majority of genes that are markers in the Sham, those that are novel markers in the T15 have not previously been associated with androgen expression or depletion, and most have no previous record of association with the prostate (Figure 5e). There was a similar pattern if we just compared the markers in the sham for each of the prostate lobes over the different timepoints. Most genes were downregulated, as expected, as these are predominantly androgen dependent, but for each lobe, there were exceptions (Figure 5d, Supplemental Figure 5c). For example, 9530003J23Rik was primarily expressed in the anterior in the Sham, though was also seen in the SV-like region, but in the T15 and T20 samples, was seen in the SV at a higher level than in the Sham and was not expressed in the Anterior (Data not shown). This was the only one of the top 20 markers for the anterior lobe shown to increase during the time course, though there were genes that were associated with the other lobes that also increased over time.

### Functional Analyses

Having determined the most differentially regulated genes following orchiectomy and that these were typically specific to each lobe and thus spatially localized within the prostate, we then wished to further understand potential functional activities these transcriptional changes could impact.

### Androgen responsiveness

As orchiectomy induces androgen deprivation, we initially focused on modulation of androgen responsive genes. AR expression was increased in both the spatial and single cell data (Figure 6a). Expression of probasin (Pbsn), a well-recognized androgen responsive gene [54] was decreased in both the single cell and spatial data (Figure 6a). Initially, it was present in the anterior, dorsal and lateral, but at lower levels in the ventral and the unknown feature. Similar to Pbsn, Msmb [55] expression, which was primarily expressed in the lateral and to a lesser extent the ventral lobes was decreased (Supplemental Figure 10a). In contrast, Fos [56] expression was upregulated. Fos was present at low levels in Sham, primarily in the lateral lobe, and at higher levels in T15, again in the lateral, though it was also seen in the dorsal at this time point. These patterns were similar in both the spatial and single cell data. Psca [57]; however was only detected in the urethra in the single cell, but across all cell types in the spatial, however, in the spatial data, it was downregulated and in most of the prostate lobes turned off (Supplemental Figure 10b).

**Figure 6.**
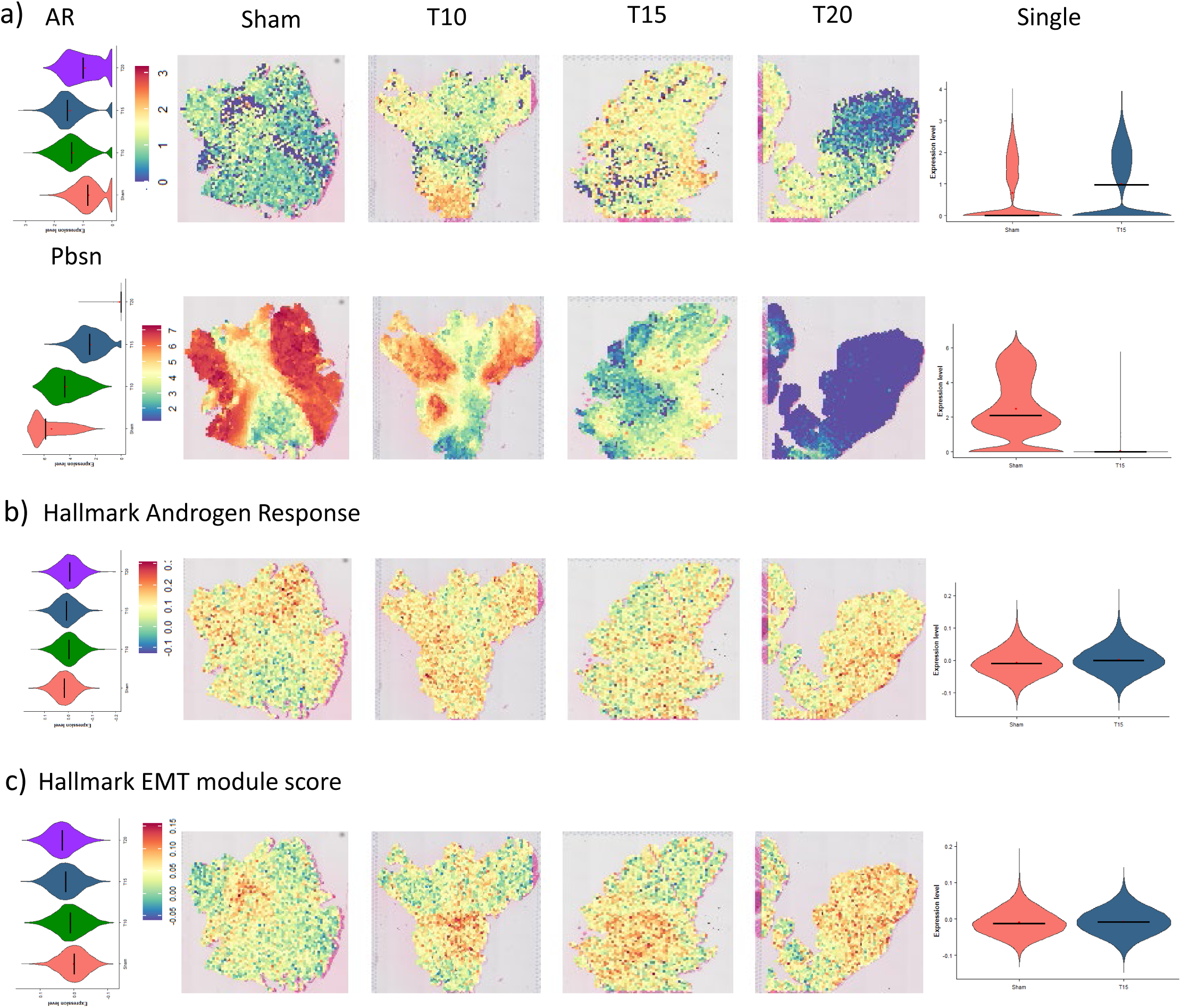
Impact of orchiectomy on androgen response. (a) Expression of genes associated with androgen response at different timepoints. Androgen receptor (AR) and Probasin are shown as violin plots and associated spatial gene expression at each time point. (b) Overall Androgen activity shown by scoring the Hallmark Androgen Response pathway using the Seurat module. (c) Hallmark EMT module score in each timepoint in left) Spatial, right) single cell spatial distribution of Hallmark EMT scores.

We also examined gene sets previously identified as being upregulated by androgen or downregulated by castration and saw a conflicting pattern with the androgen genes in the Hallmark androgen response pathway [58] decreasing over time (Figure 6b). However, analyzing specific genes that had typically been seen to be differentially expressed following surgical castration in men [59] or orchiectomy in mice [7] were generally differentially expressed in the same direction in our data, though some of them were below the level of detection and were not observed (Supplemental Figure 10c).

### Stem Cells

We examined the expression of common stem cell markers, to see if these cells were maintained following orchiectomy (Supplemental Figure 11). Again, this varied, but we saw a slight increase in Cd24a, and a larger increase in Cd44 (Supplemental Figure 11). Other reports have suggested the presence of progenitor cells, especially for luminal or basal cells [60–63] and several research groups have suggested markers for these, such as Ly6a (Sca-1) [61, 64, 65], Ly6d [66], Psca [61, 67], Runx1 [68], Tacstd2 (Trop2) [67, 69] and Zeb1 [70], but we were not able to identify such cells in our data (data not shown).

### Epithelial-mesenchymal transition (EMT)

In general, in our spatial experiment, there was an increase in mesenchymal gene expression, compared to epithelial expression (Figure 6c), however, some individual epithelial-related genes did increase, including Epcam, Cdh1, Vim and Krt8 at T15, but were lower again in the T20 (Supplemental Figure 12a,b). This increase was driven by both changes in the proportion of cells present but also changes in expression within some individual cell types, for example, there was a slight increase in Epcam expression in the anterior at T15, but this decreased again in T20 (not shown). There was also an increase in vimentin expression in T15, though in the anterior alone, this also increased in T20.

We also found a set of downregulated genes that are involved in sperm or semen production, whereas the upregulated genes were associated with a wider range of functions. Igfbp3 has previously been associated with prostate cancer [71] and both Slc40a1 (Solute Carrier Family 40 Member 1) [72] and [73] have been reported as being androgen sensitive (Supplemental Table 3), but overall, there was no consistent connection between these upregulated genes.

### Pathway Analysis

To determine pathways and functions that were altered following orchiectomy in an unbiased fashion we performed pathway analysis. We observed that pathways involved in amide and peptide binding were upregulated among others, whereas peptidase and endopeptidase activity and inhibitor activity pathways were downregulated (Figure 7a, Supplemental Figure 13). Additionally, as we might expect based on the frequency of cell types, we saw activation of immune pathways and changes in pathways involved in locomotion and motility (Figure 7a, Supplemental Figure 13).

**Figure 7.**
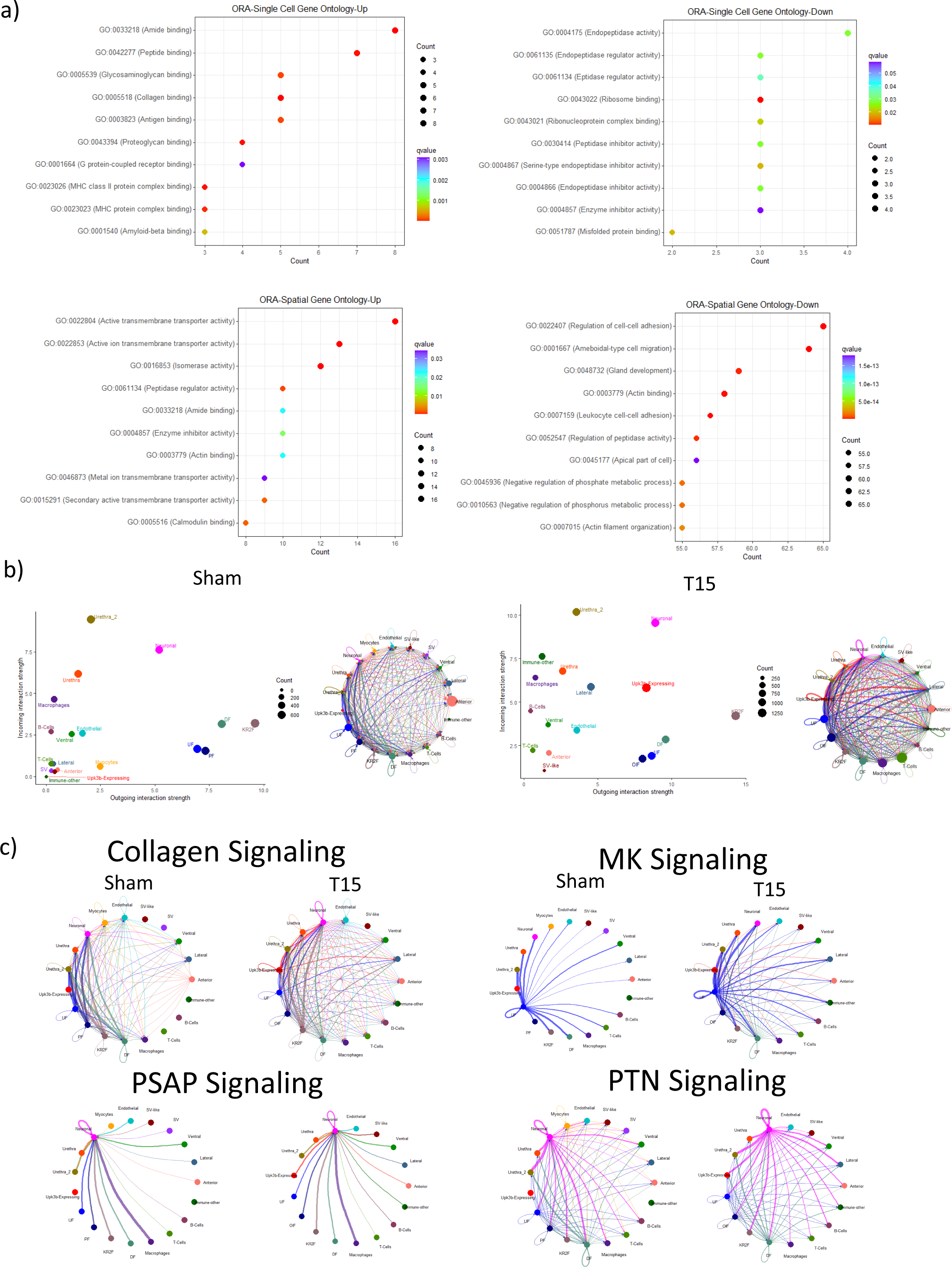
Functional analysis. (a) Gene ontology terms differentially enriched in the prostate following orchiectomy. Upper – Single cell. Lower - Spatial. Left – upregulated pathways/terms, right – downregulated pathways/terms. (b) Strength of predicted ligand-receptor interactions as calculated in CellChat in 1) Sham and 2) T15. Left, Total incoming signal vs outgoing signaling each identified cell type, Right, Total number of interactions between each cell type. (c) Signaling in selected pathways as calculated in CellChat in the single cells. Collagen, Midkine (MK), Pleiotrophin( PTN) and Prosaposin (PSAP) signaling in Sham (left) and T15 (right) cells shown.

### Ligand Receptor Interactions

Although we were able to identify pathways likely to have been differentially affected, we wanted to further investigate the mechanisms of actions by identifying specific ligand-receptor interactions that were altered in the orchiectomized mice (Supplemental 14a). Overall, the fibroblasts and the neuronal were the most active in signaling (Figure 7b). We did see a difference in the signaling patterns between the fibroblast subtypes, further supporting the idea that we had successfully identified functionally different subtypes. Consistent with changes in the pathway and gene ontology terms, and the gene expression in the different fibroblast types, we also observed expected changes in signaling in Collagen and Laminin pathways, with more cell types classed as "Influencer" for both in the Orx than the Sham (Figure 7b, Supplemental Figure 14b).

Some pathways, such as pleiotrophin (PTN) [74] and midkine (MK) [75] (Figure 7b, Supplemental Figure 14c) were primarily observed as incoming, i.e. cells were receiving signals. PTN was primarily identified in the DFs and neuronal cells, whereas MK was identified in the UFs (Figure 7c). Conversely, most cell types had prosaposin (PSAP) [76] outgoing signaling, but only the neuronal expressing cells were receivers (Figure 7b, Supplemental Figure 14d). Orchiectomy had minimal impact on PSAP signaling. MK signaling was still strongest from the UFs, but there was also signaling from the DFs and UFs, as well as the Upk3b cells (Figure 7c).

Other cell types also had specific signaling, for example, VEGF signaling was only found as outgoing signaling with the endothelial cells, and only incoming signaling with the DF, UF and KR2Fs. In general, the cells with strongest signaling in the Sham group had the strongest signaling in the Orx group, though for many of these signaling pathways, more cell types were involved in the Orx. There were some exceptions, for example (1) in Sham, macrophage signaling through TNF, which has been postulated to be involved in prostate cancer regression following castration [77] was detected in several fibroblast types and endothelial cells, but was not detected in the Orx, and others such as (2) FGF signaling [78] were significantly altered (Supplemental figure 14e). In the Sham, eight of the associated genes were detected, and activity was only predicted in the fibroblasts, most strongly the KR2F, whereas in the Orx, 16 of the associated genes were detected and the Upk3b expressing cells were also a strong source of activity in this pathway.

## Discussion

This study provides a spatiotemporal atlas of the murine prostate following orchiectomy at lobe specific resolution. Additionally, we enriched this spatial map through defining gene pathways and receptor:ligand interactions that were impacted by orchiectomy. Finally, we identified a novel fibroblast subtype, OIF, within the orchiectomized prostate.

Based on previous single cell analysis studies, each lobe of the mouse prostate has a unique expression signature [27, 28, 62]. We were able to demonstrate that previously published marker genes could be used to annotate these features in the prostate spatial data, with varying degrees of success. Even though the previous publications reported genes at the individual lobe levels, they did not look at them in a spatial context, so the marker genes could not be mapped to histological features in those studies. By viewing genes in a spatial context, we were able to identify marker genes for each of the prostate lobes and confirmed that these markers were spatially localized and their spatial relationship to each other.

Little is known about how the different mouse prostate lobes function including the specific function of the genes expressed within each of the lobes. It has been suggested that the dorso-lateral of the mouse is most similar to human prostate peripheral zone [24], though there is little consensus [18, 19]. Our results identified that the gene expression profiles of the dorsal and lateral lobes are significantly different.

While we were able to identify differentially expressed genes, the function of many of these genes were unknown in the context of prostate function and thus provide exciting opportunities to learn more regarding the functions of these genes in the context of androgen-deprivation in the prostate and how these changes may impact prostate cancer. Orx induced differential expression of several hundred genes. Many of these had previously been identified as (1) being expressed in the prostate and/or androgen regulated [7, 79], (2) as promoting [80] or suppressing [81] prostate cancer progression and (3) were often localized to specific prostate lobes [24, 27, 28]. Downregulation of the majority of gene expression occurred rapidly, by day 10, though a portion of genes showed were downregulated at later time points. Upregulation of some genes showed a similar temporal pattern of altered expression. As 10 days post-Orx was the first timepoint evaluated, it is possible that gene expression changes occurred even earlier than this period.

The temporal nature of our study and ability to identify lobe-specific marker genes allowed us to determine how the prostate changed over time following Orx. Interestingly, this was also the case for the urethra including the muscular part of it. This was most notable at day 20, when Actg2 was strongly expressed in the muscle. This was also the case with Acta2 and Actb, which were strongly expressed in the muscle on day 20, but much weaker on the earlier timepoints. Curiously, on these earlier timepoints, these actins were not strongly associated with the urethra muscle, and this feature even had the lowest expression. There were also marked gene expression changes in the fibroblasts. However, immune cells, and endothelial cells showed minimal differentially expressed genes as demonstrated on the T15 UMAP where the sham cells overlapped each other.

As Visium data is generated at resolution of 50 μm spots, each spot consists of multiple cells (typically between 4 to 10), thus in the current study, we mapped single cell data derived from prostate single cell suspensions to the Visium spatial data to enhance the resolution that the Visium-generated gene expression provided. This methodology was very effective and provided a window into the types of cells and Orx-induced changes in the different lobes.

We looked at the expression of cytokeratins that are markers for basal or luminal cells [82]. Krt5 and Krt 14, both basal markers were detected at very low levels in the single cell data. They were detected more in our spatial data, with Krt14 especially increasing over time, with the largest increase in the anterior (Not Shown). However, Krt8, commonly a luminal marker, showed the same pattern of higher expression in the Orx.

Additionally, other putative luminal markers such as Psca or Wfdc2 were primarily detected in the urethra and ventral lobe, which may indicate the ventral lobe has increased stemness. However, as the levels of these were low, and few cells expressed the luminal genes but not the basal genes, it was difficult to use these to label any cells as either basal or luminal. Previous reports had identified sub populations of basal and luminal cells, many of which were either castration resistant, or could act as progenitor cells [62, 69, 83], however, we did not confidently identify populations of any of these cells based on the proposed marker genes.

In addition to changes in epithelial cells, we identified Orx-induced changes in non-epithelial cells. In terms of fibroblasts, our study identified 4 fibroblast subtypes in both Sham and Orx prostates. The gene expression patterns identified them as ductal fibroblasts/R1/subglandular fibroblasts, as previously identified [28, 31, 32], prostate and urethral fibroblasts and the R2/Interstitial fibroblasts [28, 32]. Fibroblast markers identified in a previous study [27] poorly differentiated between these and the prostate fibroblasts and it may be that other groups have combined these two into one subset. Kwon et al. [32] identified an R3 fibroblast, which were likely myofibroblasts, and were not detected in our study. We did see a change in the type of fibroblasts. The R2 showed a significant increase in frequency. As these had also been identified as interstitial fibroblasts, this may explain the increase in detection.

We also identified a novel subset of fibroblasts in the Orx that we named orchiectomy-induced fibroblasts (OIF). These made up a similar proportion of cells 14% of total cells, and 32% of the fibroblasts in the Orx as compared to the prostate fibroblasts, which made up 11% of all cells and 28% of the fibroblasts in the sham. This suggested that the OIFs may have been derived from the prostate fibroblasts.

However, they did not demonstrate strong expression of the prostate fibroblast marker genes as identified by us or Joseph et al. [31]. This finding is in contrast with the other fibroblast subtypes we observed in that although many Orx-induced changes in expression could be identified, they could still be identified as their respective fibroblast subtype-based expression of their marker genes. We did find some similarities between OIF and prostate fibroblasts in terms of their ligand:receptor interactions and signaling pathways. Specifically, despite there being several different pathways identified between OIF and prostate fibroblasts, the FGF and IGF pathways were among the most represented pathways in both of the fibroblast subtypes.

Additionally, pathway analysis indicated that fibroblasts subjected to androgen deprivation had altered expression of collagen types and other extracellular components. This phenomenon could provide a mechanism to account for the observed phenotype, in which the post-Orx prostate is less organized and has greater proportions of extracellular matrix.

Immune cells were the other major cell type that were represented in the non-epithelial cell population. Unlike fibroblasts, we did not detect differences in the types of immune cells in the Orx, but did observe both a change in the overall percentage of immune cells in the total cell population and a change in the relative distribution of each cell type. Overall, about 10% and 36% of the total cells were immune cells in the prostates from Sham versus Orx groups, respectively. It was unclear if this change was due to Orx-induced increase of immune cell recruitment due to a relative change secondary to an overall decrease of the prostate epithelial cell population in the Orx-group prostates.

In addition to the novel OIF cell, we identified another potentially novel cell type in the prostate, termed Upk3b expressing. Though the Upk3b cells were distinct from urethral cells, both on the UMAP and in expression of the top marker genes, we considered that they were potentially urethral endothelial or mesothelial. These showed differences between Sham and Orx, for example in Midkine signaling, though this may have been due to the low number of these cells in our Sham cells.

There are several limitations of the current study that should be recognized. For the spatial data we were able to use the histology to confirm that the clusters actually represented their associated tissues; however, we were not able to do so for the single cell data. Separating the lobes before running single cell would give us greater confidence that they had been identified correctly. While this would have been relatively simple for the Sham group, it would be very challenging for the Orx due to the marked involution of the lobes. We leveraged data from other publications in which lobe specific single cell analysis was performed to perform cell annotation [24, 27, 28]. We had validated marker genes for the prostate lobes in our spatial data which were consistent with previously published single cell and microarray data, however the fraction of cells comprising the lateral and ventral lobes were low. As we felt confident in our markers and our identification of lobes with these genes clustering separately on the UMAP, we felt it likely we have accurately identified these features and that the low percentage was likely due to a differential effect of Orx-induced involution of the different lobes. However, we cannot rule out that certain cell types were selectively lost when the single cell preparation was made.

In conclusion, our study provides a spatiotemporal atlas of Orx-induced changes in the mouse prostate. Furthermore, a novel Orx-induced fibroblast was identified. Taken together, these findings provide an enhanced understanding of the Orx-induced changes in the prostate that can inform both murine models of prostate cancer and androgen biology.

## Acknowledgements

Research reported in this publication was supported by the National Cancer Institute under Award Number P30 CA046592 by the use of the following Cancer Center Shared Resource: Single Cell Spatial Analysis Shared Resource. This project was funded, in part, by the Richard and Susan Rogel Oncology Professorship at the University of Michigan Rogel Cancer Center.

## Data availability

Data are available on the Gene Expression Omnibus (GEO) as follows: Day 0 and Day 15 Visium data are available as GSE284571; Day 10 and Day 20 Visium data are available as GSE295043; and Day 0 and 15 single cell RNAseq data are available as GSE295192.

## Notes

https://www.ncbi.nlm.nih.gov/geo/query/acc.cgi?acc=GSE284571

### Competing Interest Statement

The authors have declared no competing interest.

