## Supplemental Figures and Legends for "A spatiotemporal atlas of orchiectomy-induced androgen deprivation-mediated modulation of cellular composition and gene expression in the mouse prostate"

#### Supplemental Figure 1 - Outline of experimental plan

Mice were sham orchiectomized (Sham) or orchiectomized and prostates harvested for spatial transcriptomics at Sham, 10 (T10), 15 (T15) or 20 (T20) days later. Additionally, some prostates were taken at Sham and T15 for single cell analysis.

#### Supplemental Figure 2 - Tissue Sections for Analysis

a) Placement of the sample on the Visium slide. Prostate section was placed on the visium slide, with varying proportions of the sections in the analyzed area.
b) QC metrics for each sample, colored by timepoint. 1) Number of spots per section for each of the timepoints. 2) Number of genes detected in each sample, 3) Number of genes per UMI for each sample.
c) UMAPs of replicate samples from each time point.
d) Sham prostate annotated by clusters identified in Seurat.
e) Correlation matrix for Pearson coefficient of each sample for the four timepoints as calculated in the corrplot package.
f) Correlation matrix for Pearson coefficient for the four timepoints

#### Supplemental Figure 3 - Specific sections

For each timepoint, one sample was chosen that had the most spots for the primary visualization.

a) Distribution of counts and genes spatially across the four timepoints. Top counts per spot, bottom, genes per spot.
b) Violin plots of number of counts (top) and genes (bottom) on each timepoint. White bar- median, Red dot – mean.
c) Each sample was analyzed without the presence of the other time points. UMAP, colored by cell type assigned to the spot. Each cluster is circled to demonstrate the spatial separation on the UMAP.

#### Supplemental Figure 4 - Annotation using previously published data

Module score of each spot using differentially expressed genes for each lobe identified in 1) This paper, 2) Berquin, 3) Graham, 4) Crowley. Columns show annotation based on 1) Anterior, 2) Dorsal, 3) Lateral, 4) Ventral.

#### Supplemental Figure 5 - Analysis of Anterior lobe

a) Markers for clusters primarily seen in the Sham anterior.
b) Anterior only, Sham spots colored by Sham only clusters. Inset initial clusters.

c) Expression of selected marker genes for the clusters identified in the Anterior lobe over time. Marker expressed in at least 50% of the cluster. Top 2 genes for each cluster shown. Expression of the genes in the anterior shown as a feature plot on the UMAP and in the Sham as spatial.

#### **Supplemental Figure 6 - Analysis of Dorsal lobe**

a) Dorsal spots from each timepoint combined into a separate object and re-analyzed, UMAP, colored by i) Cluster, ii) Timepoint
b) Dorsal spots from the Sham sample only taken and re-analyzed.
c) Expression of top two markers for each cluster identified in the Sham dorsal, top row, dorsal across all timepoints, second row dorsal Sham only
d) Spatial expression of these markers

#### **Supplemental Figure 7 - Immune gene expression**

Expression of further selected immune genes

a) Cd74.Left spatial distribution. Right Violin plots of Spatial (upper) and Single Cell (lower).
b) Expression of Cd14 in single cells.
c) Expression of Cd3d in single cells.
d) Estimated distribution of macrophages in the Sham as predicted by Top) Card deconvolution and Middle) Seurat module score from the top 100 differentially expressed genes. Bottom) Macrophage module score for each identified cell type in the single cell.
e) Expression of selected genes in Sham and T15 spatial and Single cell (Violin plot) i) Prprc, ii) Adgre, iii) Cd3d,, iv) Cd19, v) Cd79a.

#### **Supplemental Figure 8 - Temporarily regulated genes**

a) Selected genes showing different patterns of change over time – Top row) Svs5, Actg2, Second row) Clu, Pate7, Third row) 9530002B09Rik, Nupr1

b) Selected genes specific to each lobe. Top row) Anterior – Gsdma, Ren1, Second row) Dorsal – Nlpr1-, Pate14, Third row) Lateral - Smgc, Sp5, Fourth row) Ventral – Nxpe4, Sbp

#### **Supplemental Figure 9 - Deconvolution**

a) Deconvolution of different cell types detected in the single in the spatial data. Module score was calculated based on the most differentially expressed genes for each cell type in the single cell data.

#### **Supplemental Figure 10 - Androgen responsive genes**

Expression patterns of selected androgen responsive genes.

a) Expression plots of downregulated genes Left Single cell violin plots, Right Spatial, 1) Violin plot, 2) Spatial ExpresioRow1 Pbsn, Row2 Msmb, Row 3 Fos, Row4, Psca.

b) Expression plots of upregulated genes Left :Single cell violin plots; Right: Spatial data.

c) Expression of genes identified as androgen regulated in previous published work. Expression of genes identified as androgen regulated in previous published work. Top) Following castration in human (Vaarala,2012), Bottom) Following castration in mouse (Wang 2007). Left) Single cell violin plots; Right) Spatial.

*Vaarala, M. H., et al. (2012). "Identification of androgen-regulated genes in human prostate." Mol Med Rep 6(3): 466-472.*

*Wang, X.-D., et al. (2007). "Expression profiling of the mouse prostate after castration and hormone replacement: implication of H-cadherin in prostate tumorigenesis." Differentiation 75(3): 219-234.*

#### **Supplemental Figure 11 - Expression of other selected genes**

Expression of selected stem cell markers, Cd24a (top) and Cd44 (bottom) in spatial (left) and single cell (right) data.

#### **Supplemental Figure 12 - Expression of epithelial and mesenchymal marker genes**

a) Selected epithelial (Epcam, Cdh1 and Krt8) marker expression in spatial (top and middle) and single cell (bottom), Spatial shown as overall all expression for each timepoint, then as expression in each region for Sham and T15. Single shown as expression in each cell type

b) Selected mesenchymal (Vim, Acta2 and Fn1) marker expression in spatial (top and middle) and single cell (bottom), Spatial shown as overall all expression for each timepoint, then as expression in each region for Sham and T15. Single shown as expression in each cell type

#### **Supplemental Figure 13 - Gene Ontology and Pathway Analysis**

a) Pathways enrichment in single cell from multiple datasets. Left) Enriched in Sham, Right) Enriched in T15.
b) KEGG Pathways enrichment in single cell. Left) Enriched in Sham, Right) Enriched in T15.

#### **Supplemental Figure 14 - Ligand-Receptor Interactions**

a) Overall Ligand-Receptor interactions in left) Sham and right ) T15 calculated in single cell data.

i. Heat map of the top 20 pathways identified in the Sham single cell showing left) outgoing (i.e., signaling from a cell) and right (i.e., signaling to a cell).

ii. Overall interactions by left) predicted number of interactions between two cell types and right) predicted strength of interactions between two cell types.

iii. Scatter plot of predicted strength of interactions in each cell type showing the outgoing strength and incoming strength for each cell type.

iv. Heat map and river plot of interactions, top) incoming, bottom) outgoing. The patterns reveal how the cells coordinate with each other as well as how they coordinate with certain signaling pathways to drive communication with hierarchical clustering of cells and pathways. Left) heat map, right) River plot.

b-e) Selected pathways for Col1) Sham, Col2) T15, showing:

b) Left) Collagen, Right) Laminin.
c) Left) PTN, Right) MK.
d) Left) PSAP, Right) ADGRE.
e) Left) TNF, Right) FGF.

i. Signaling roles for selected pathway for each cell type.

ii. Heat map of strength of interaction between each cell type for selected pathway.

iii. Violin plots for expressed genes in each of the pathways

iv. Overall interactions by predicted number of interactions for selected pathway between two cells.

v. Scatter plot of predicted strength of interactions for selected pathway in each cell type showing the outgoing strength and incoming strength for each cell type.

### Supplemental Figure 1: Outline of experimental plan

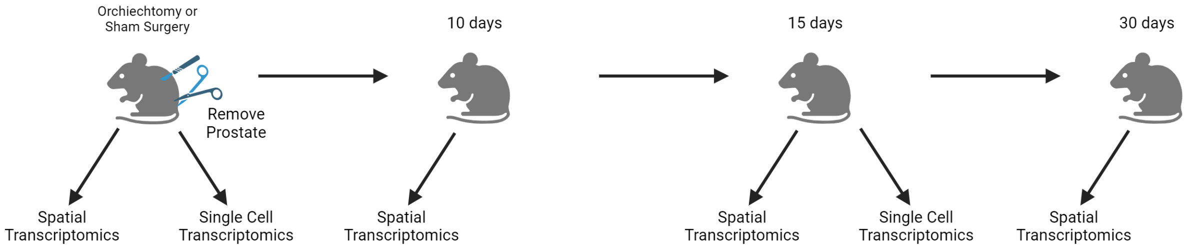

### Supplemental Figure 2

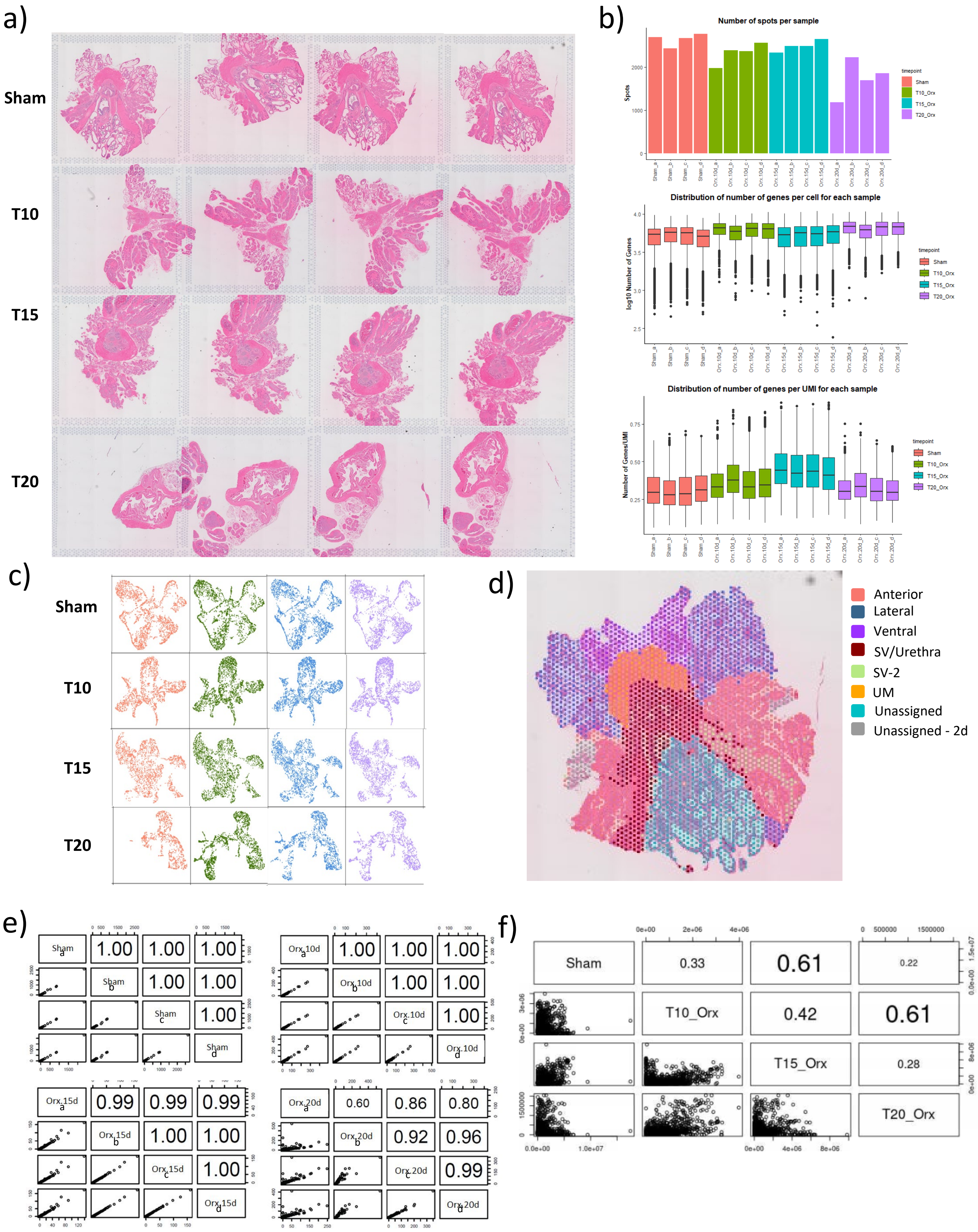

### Supplemental Figure 3

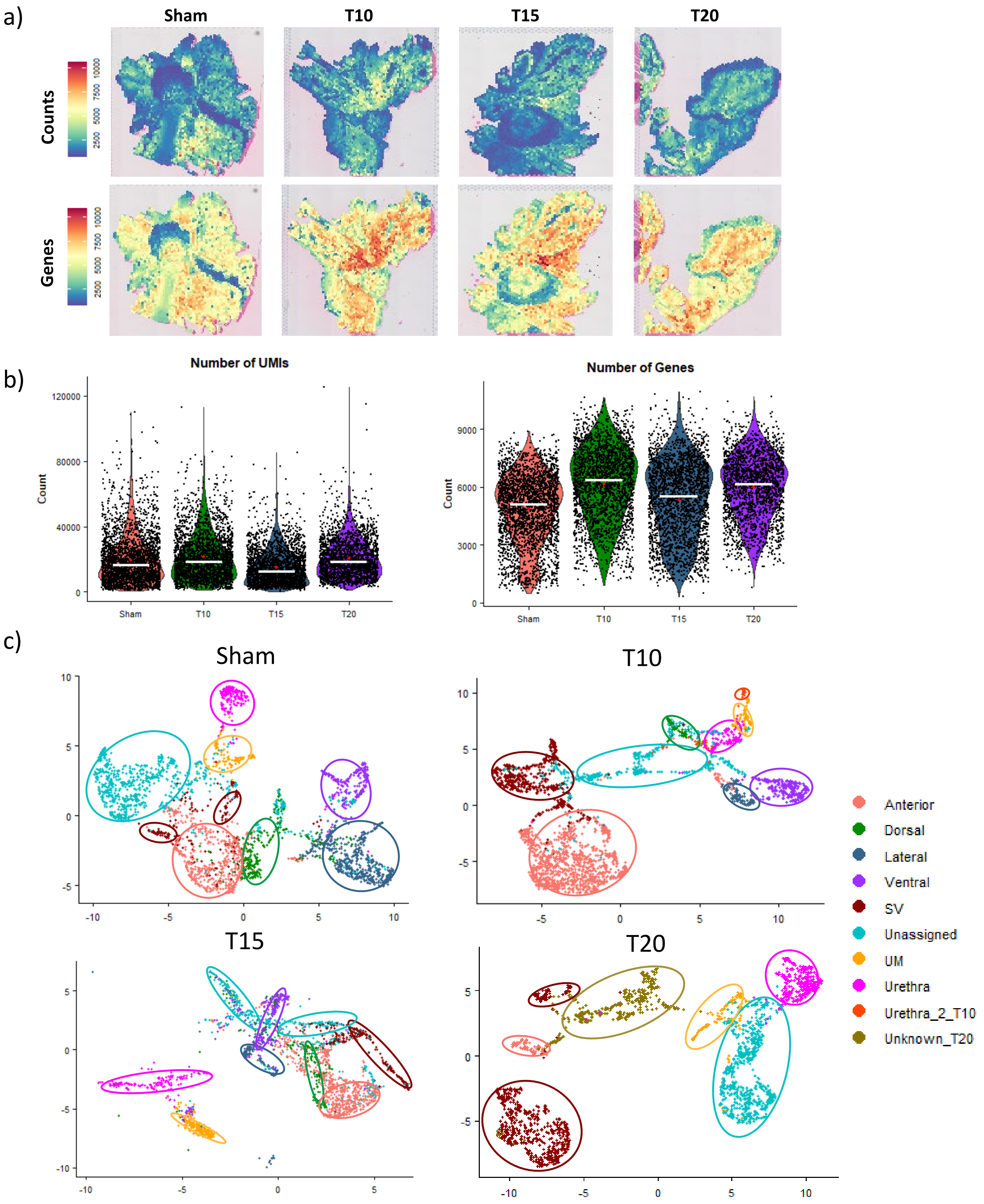

### Supplemental Figure 4

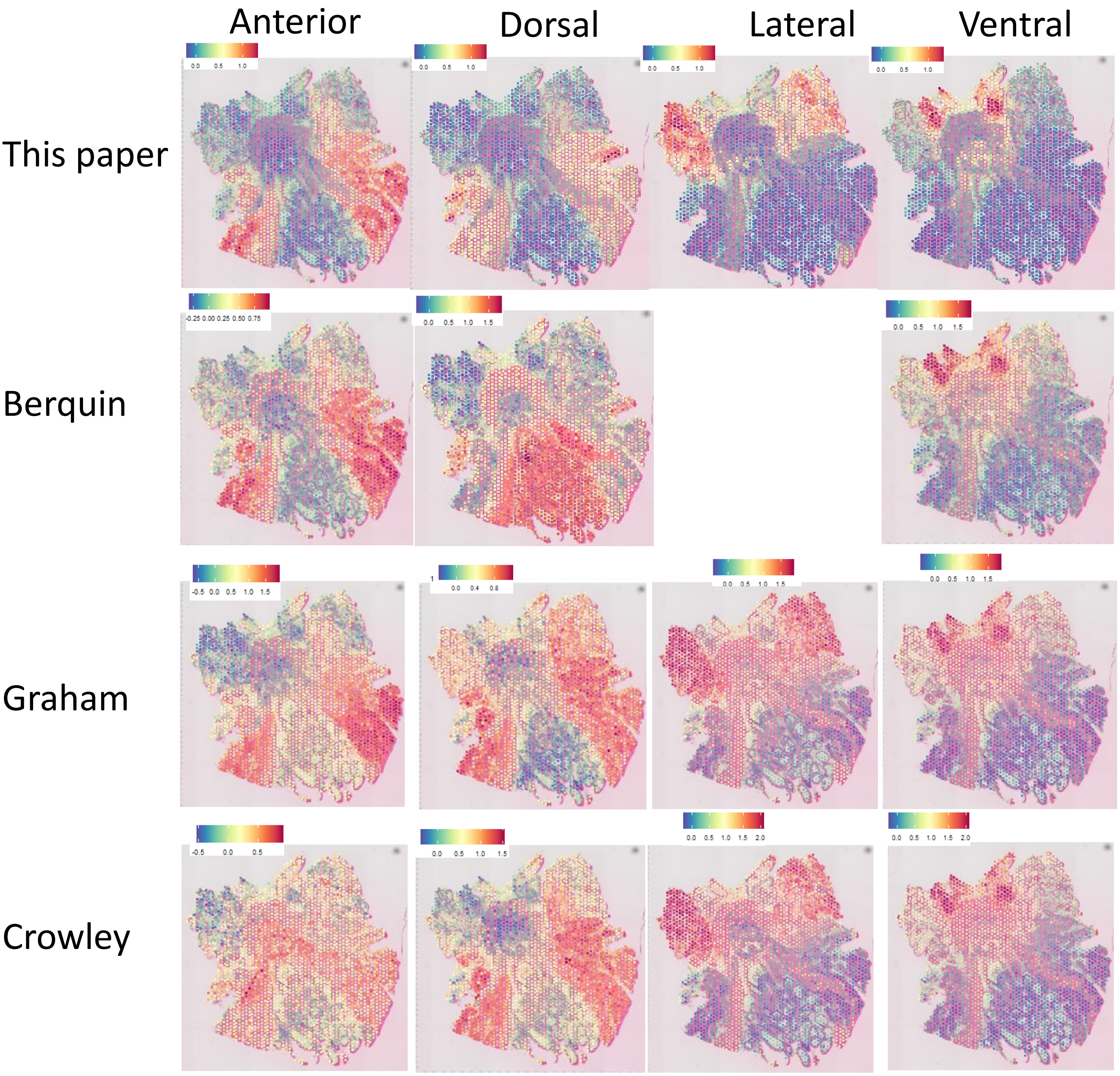

### Supplemental Figure 5

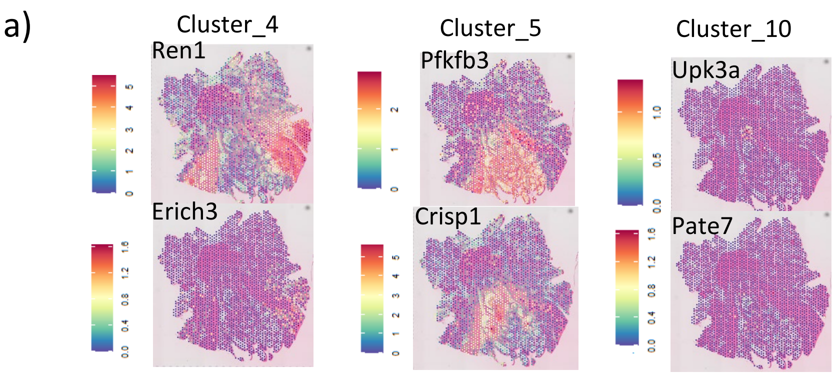


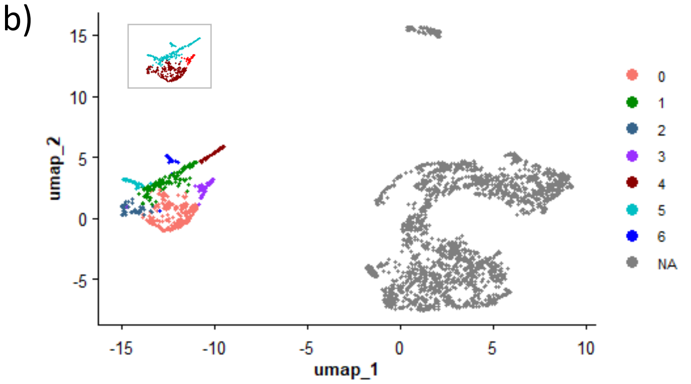


| Cluster | Spots |
|---|---|
| 0 | 200 |
| 1 | 118 |
| 2 | 50 |
| 3 | 45 |
| 4 | 44 |
| 5 | 29 |
| 6 | 24 |

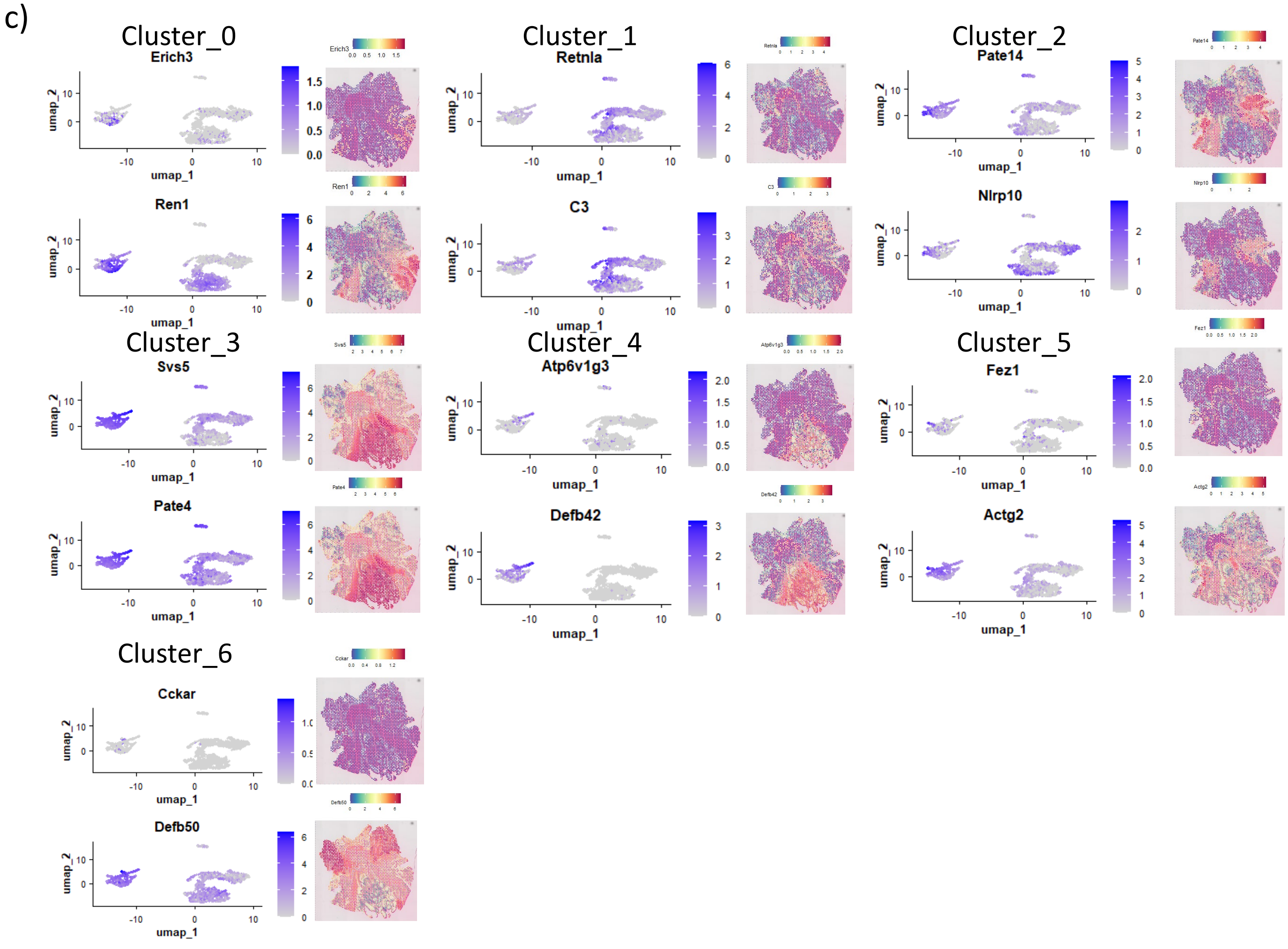

### Supplemental Figure 6

ai)

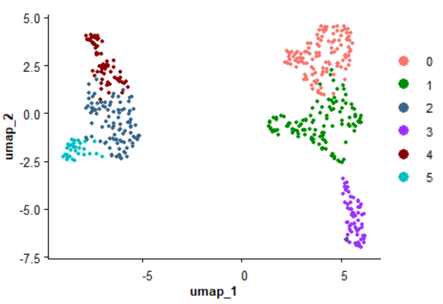


aii)

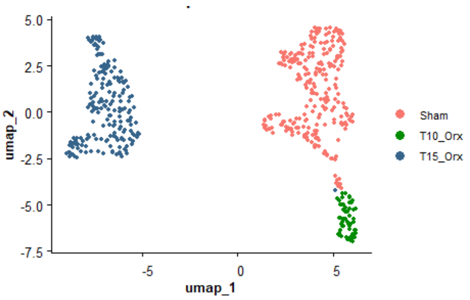


b)

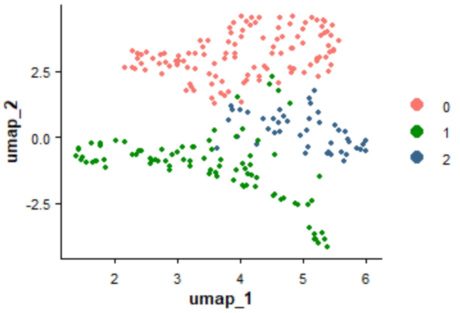


| Cluster | Spots |
|---|---|
| 0 | 116 |
| 1 | 85 |
| 2 | 45 |

c) Cluster_0 Cluster_1 Cluster_2

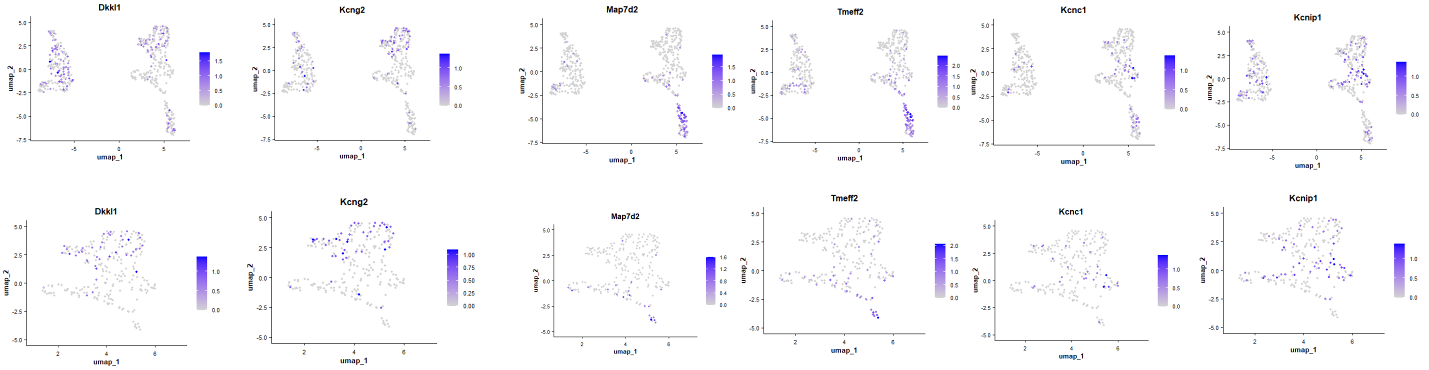


d)

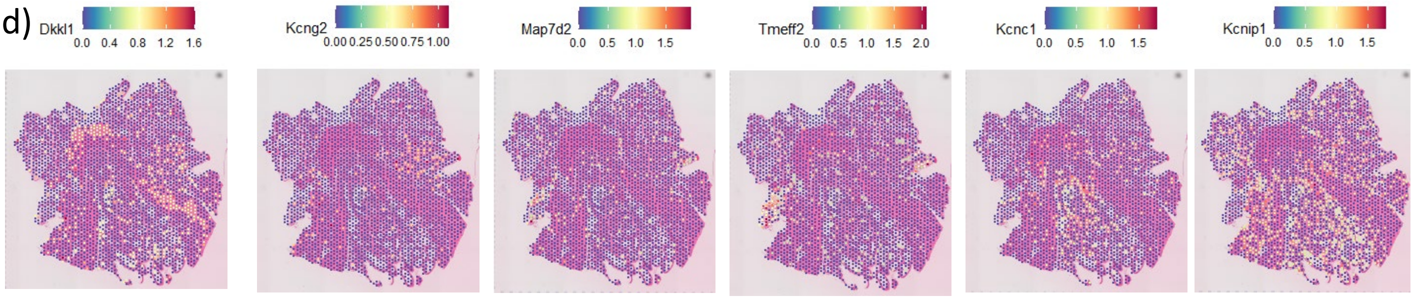

### Supplemental Figure 7

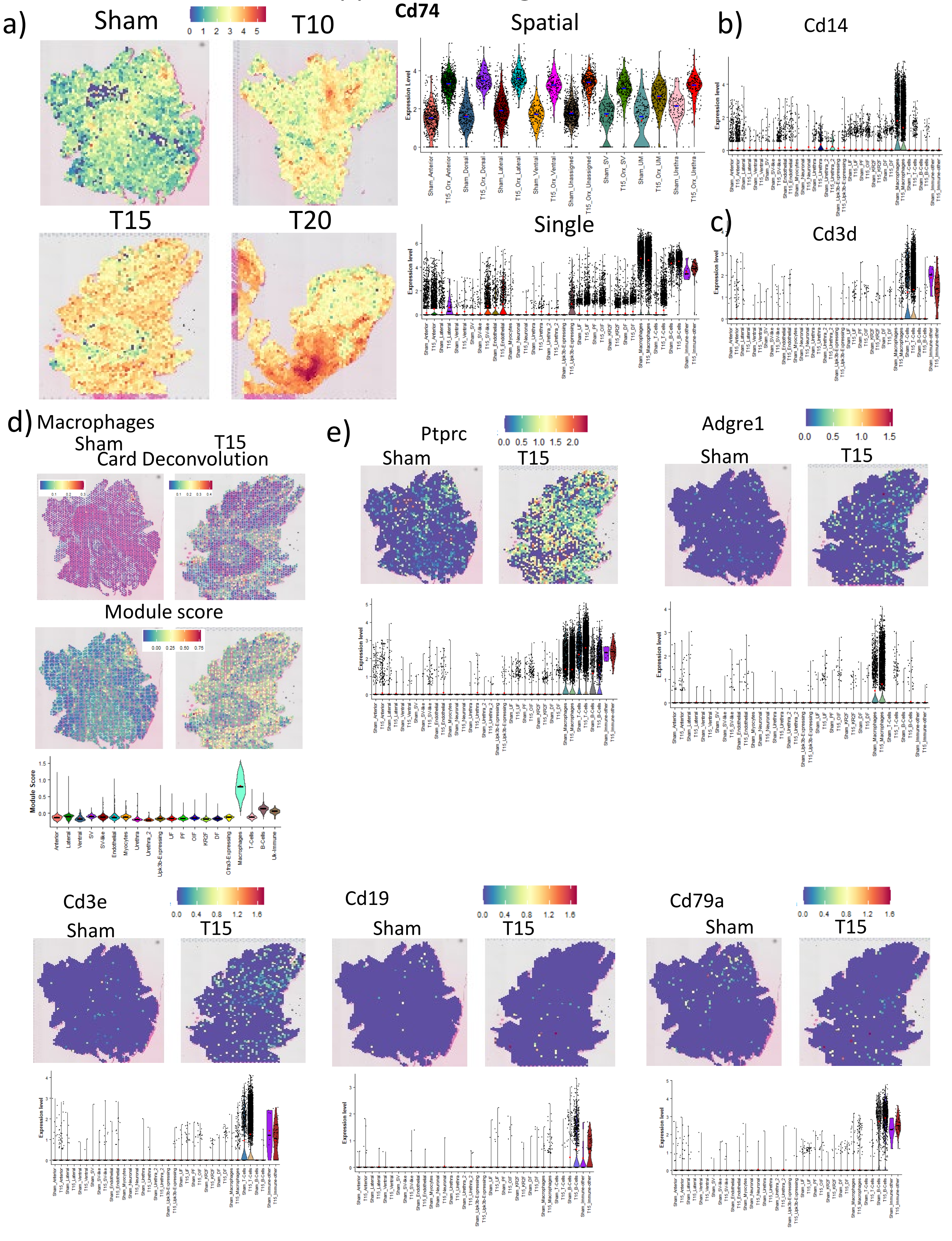

### Supplemental Figure 8

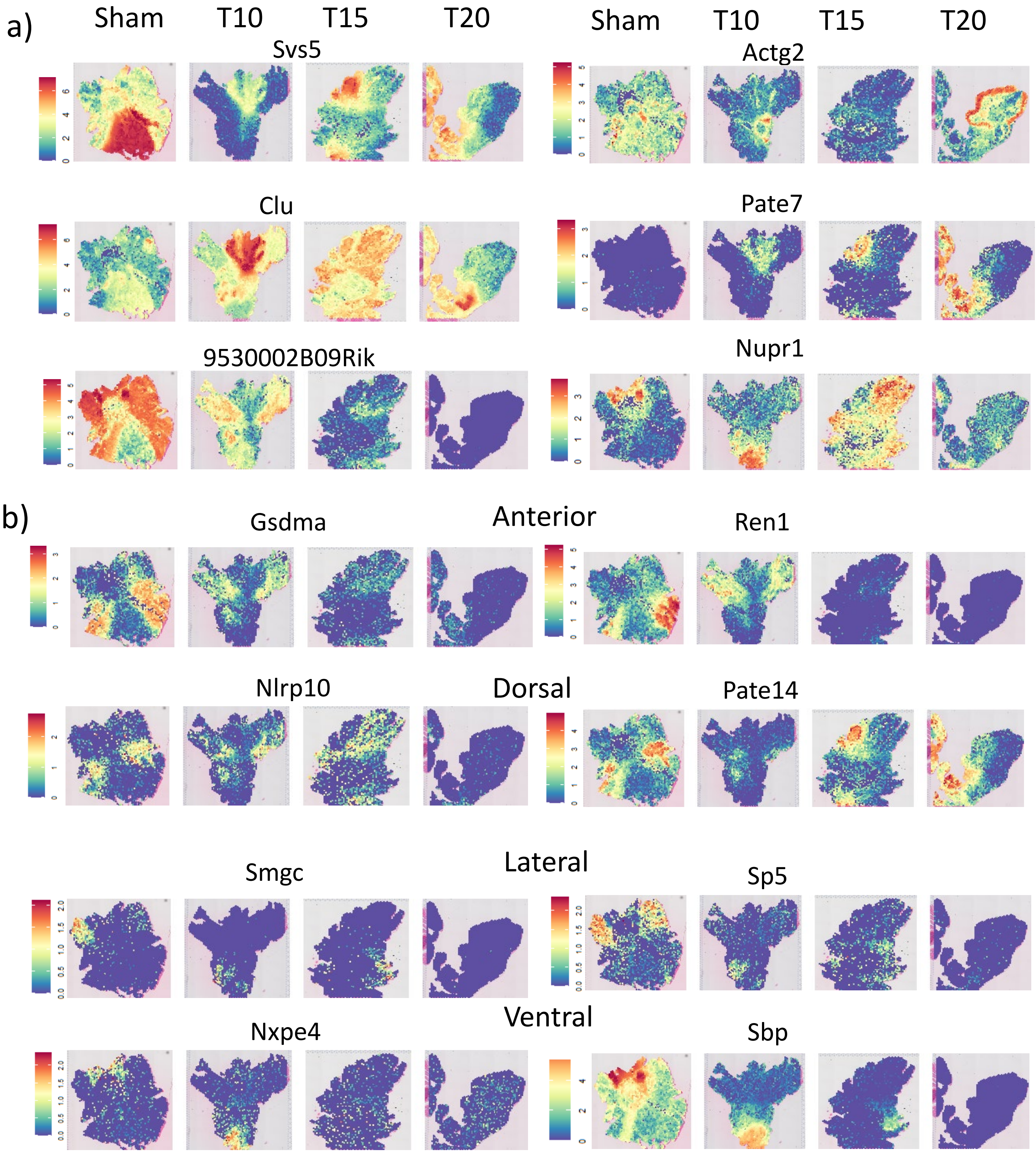

### Supplemental Figure 9

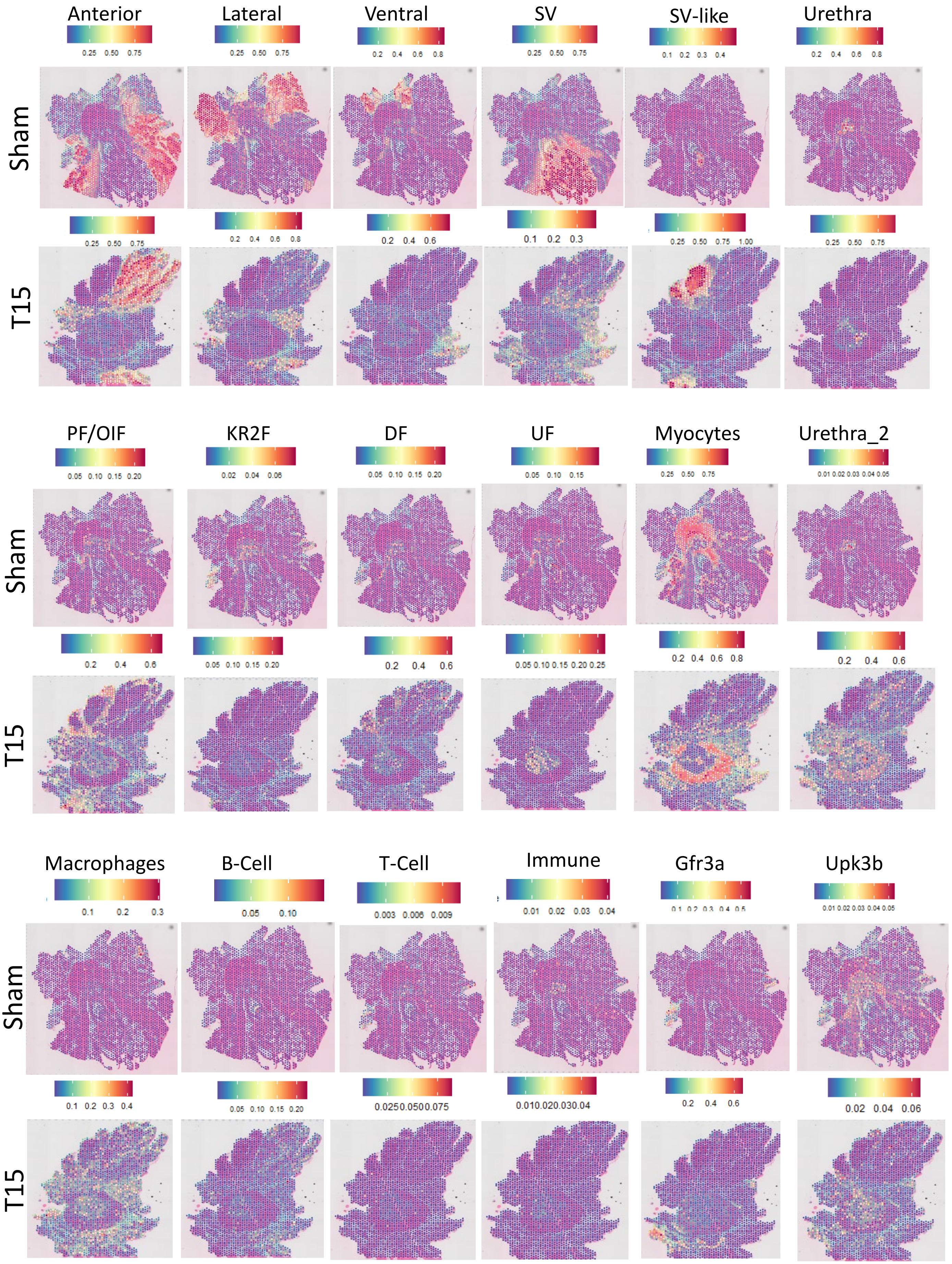

### Supplemental Figure 10

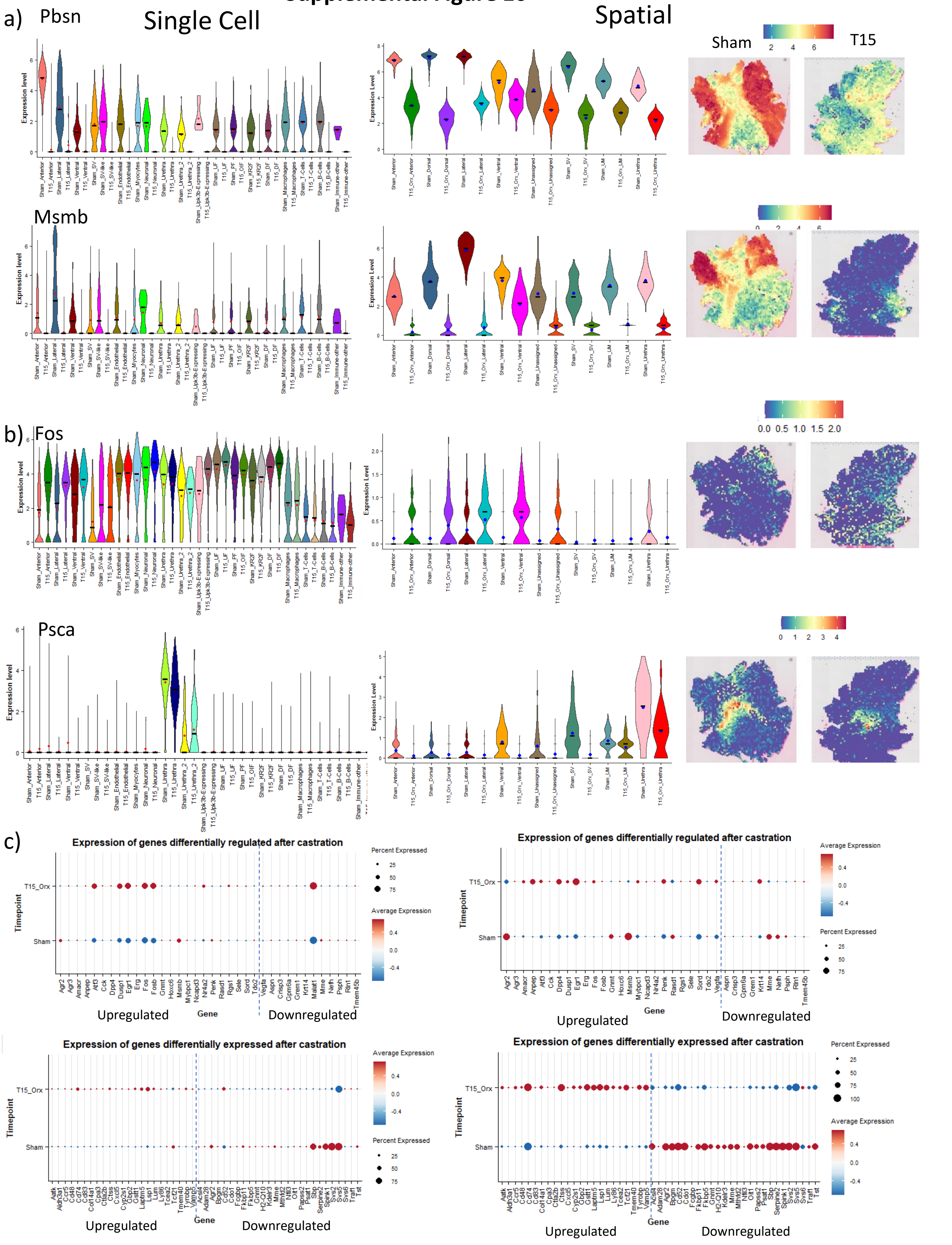

### Supplemental Figure 11

## Cd24a

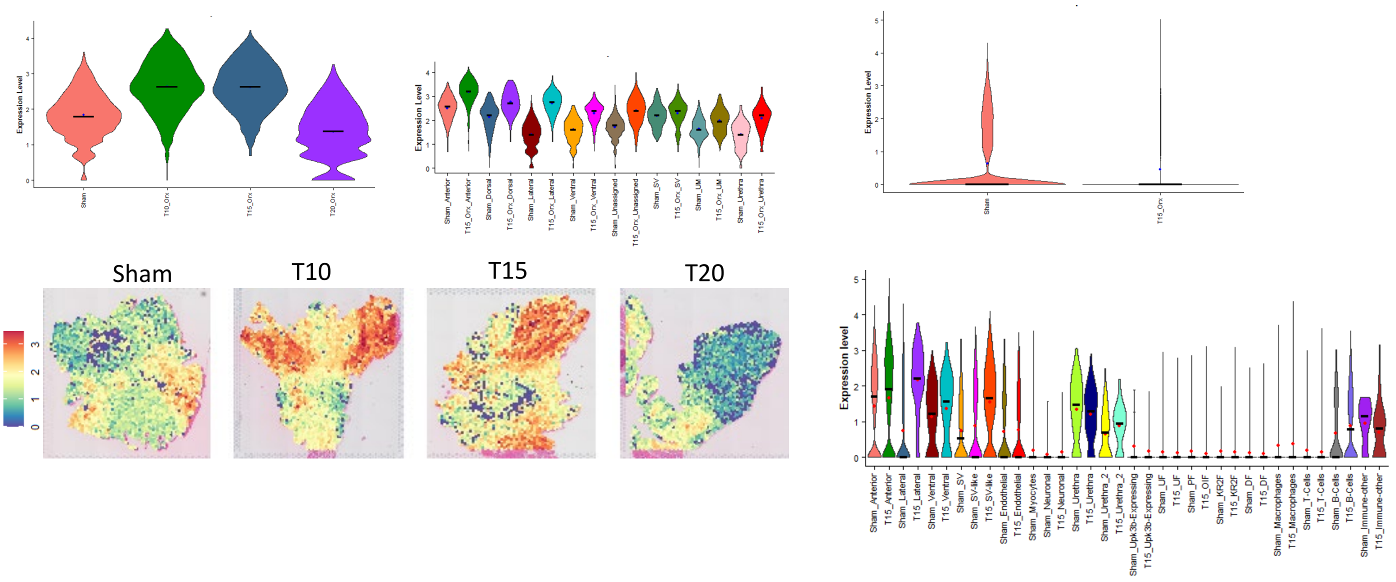


## Cd44

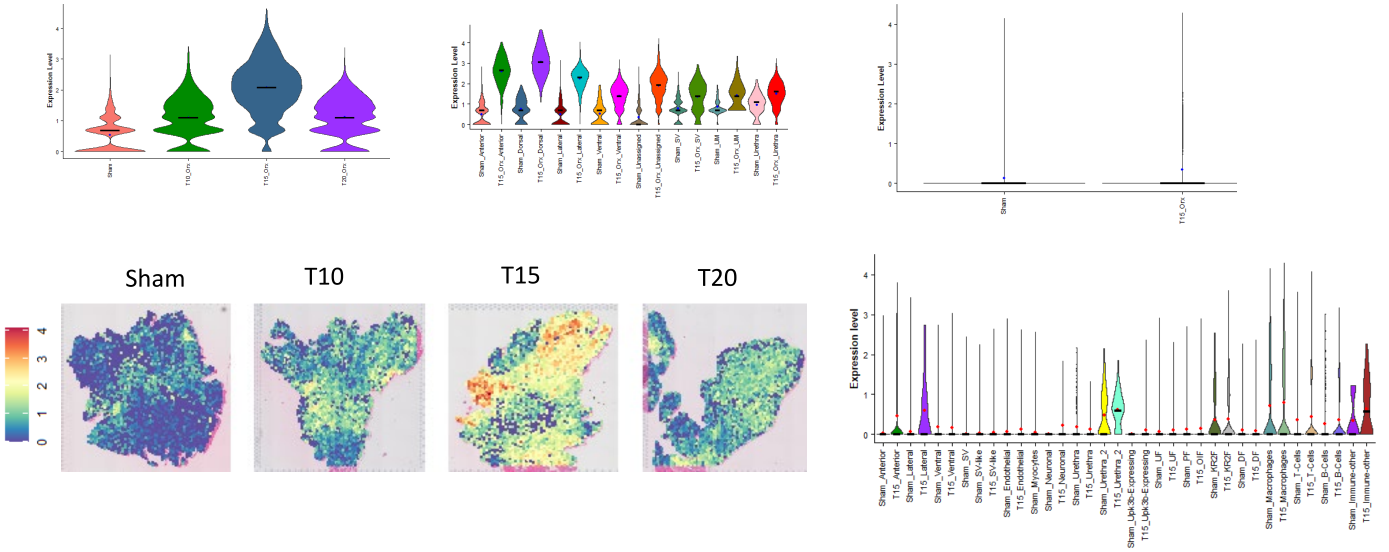

### Supplemental Figure 12

a)

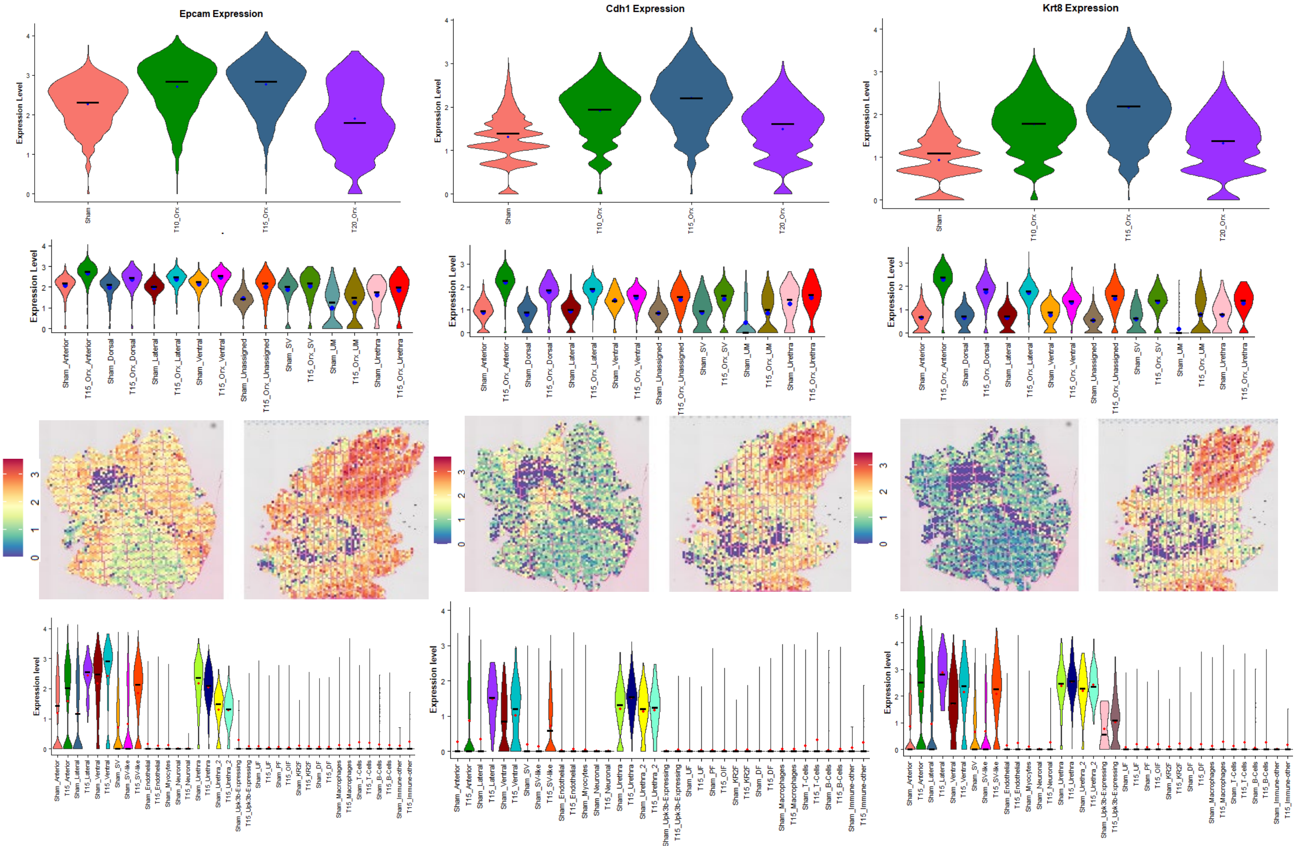


b)

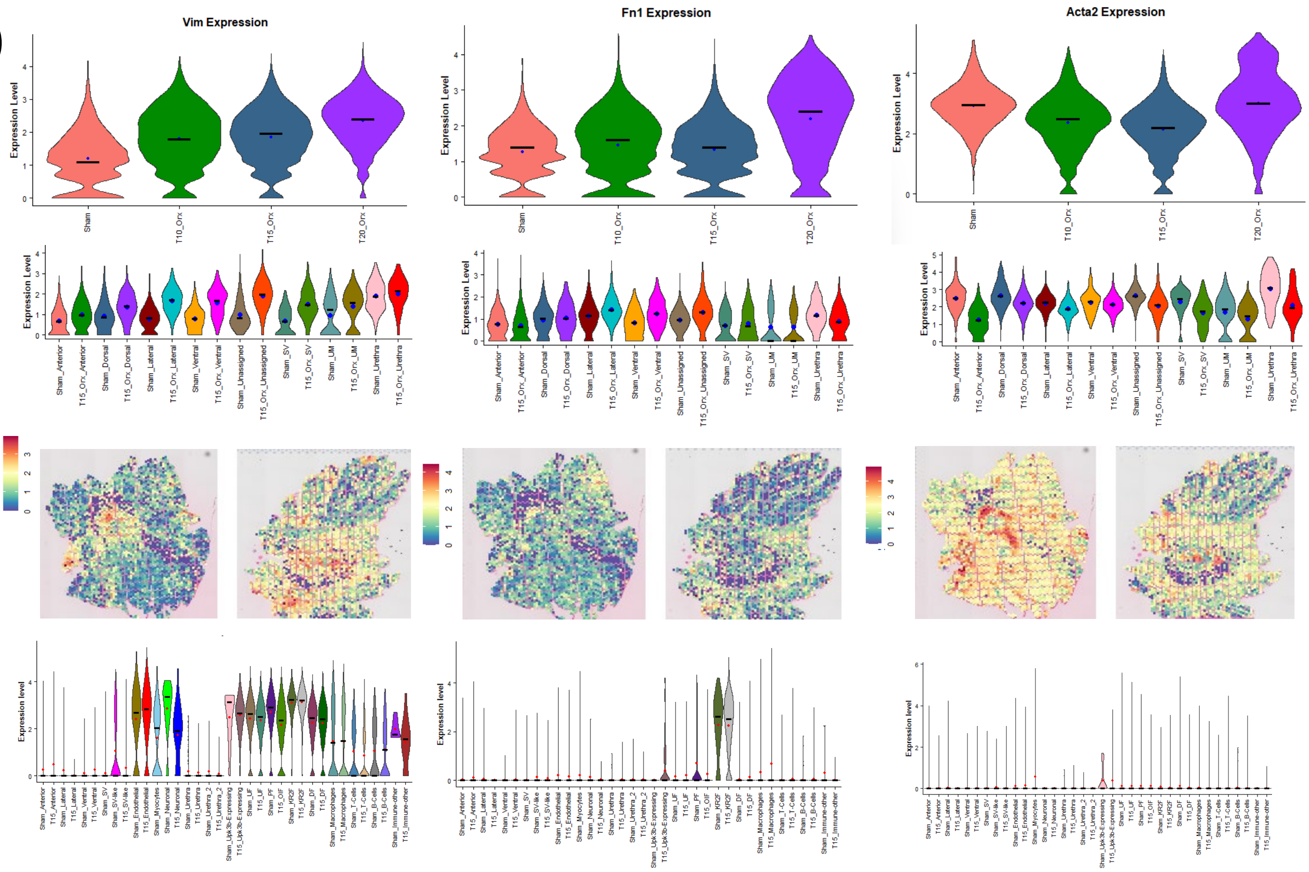

### Supplemental Figure 13

#### A. Combined pathways

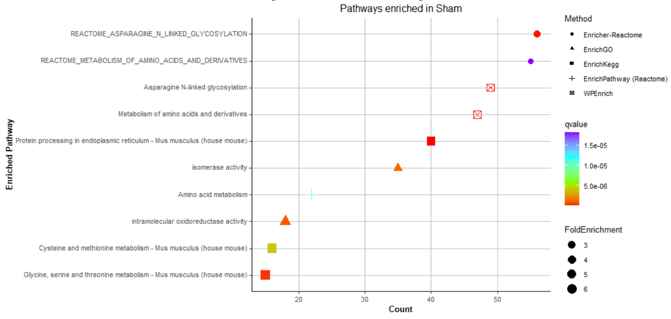


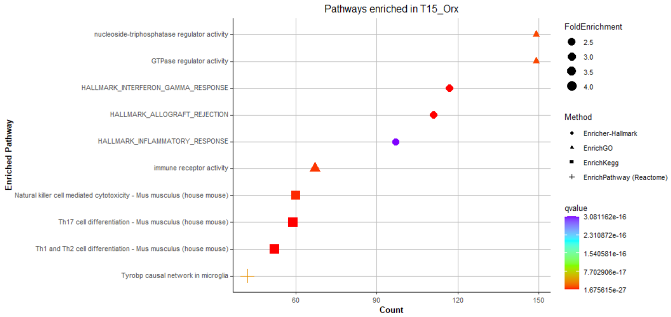


#### B. KEGG pathways

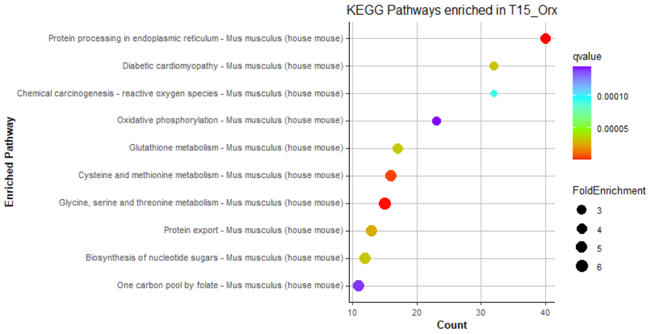


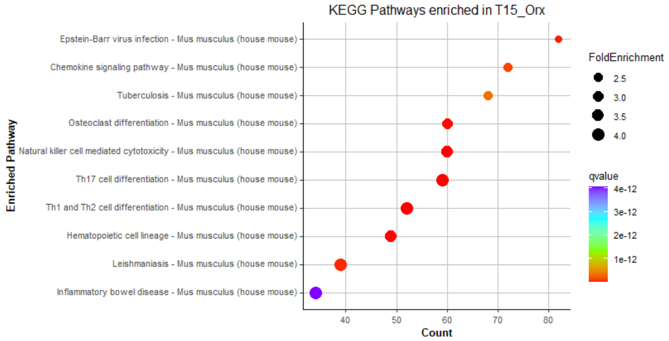

### Supplemental Figure 14a

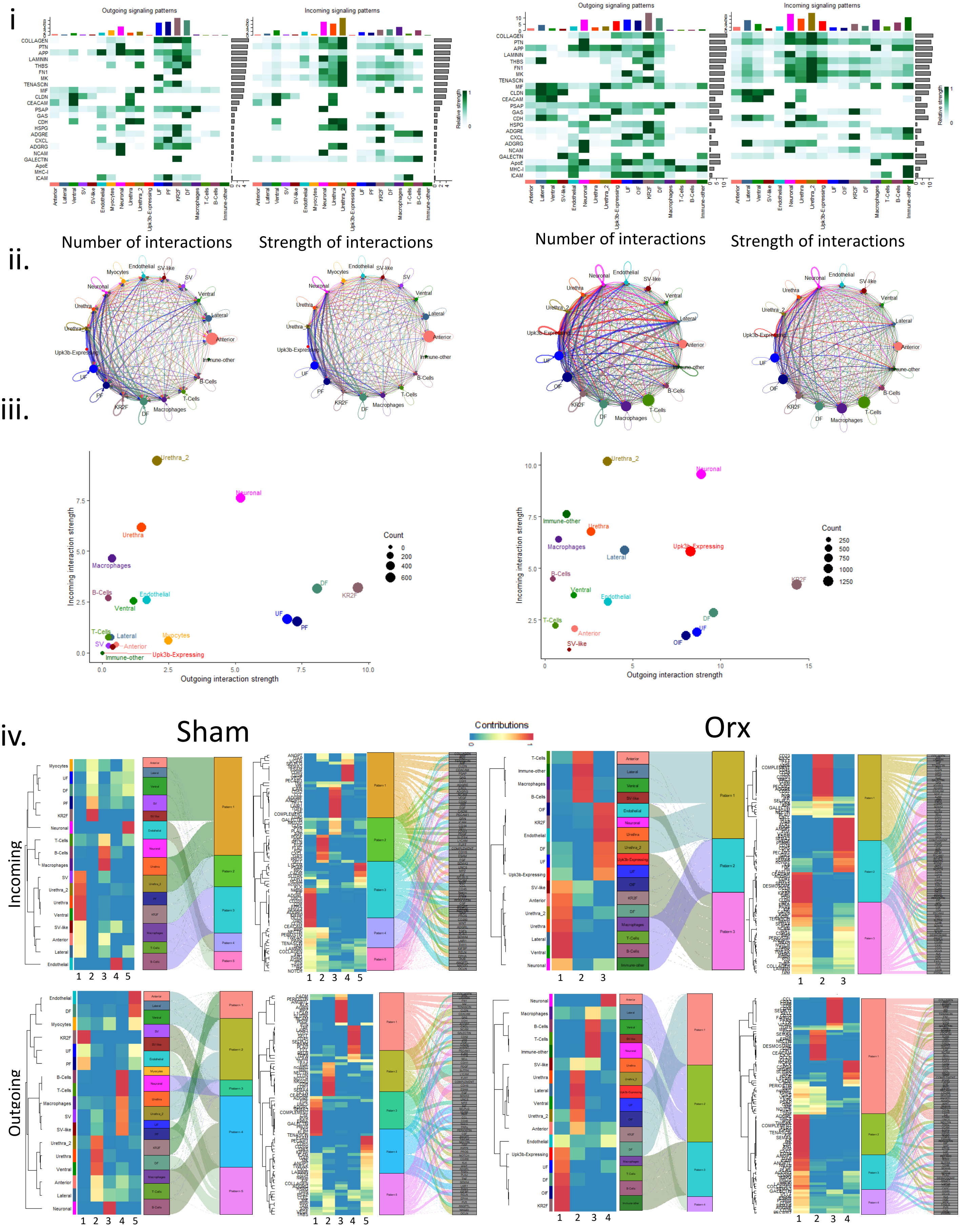

### Supplemental Figure 14b

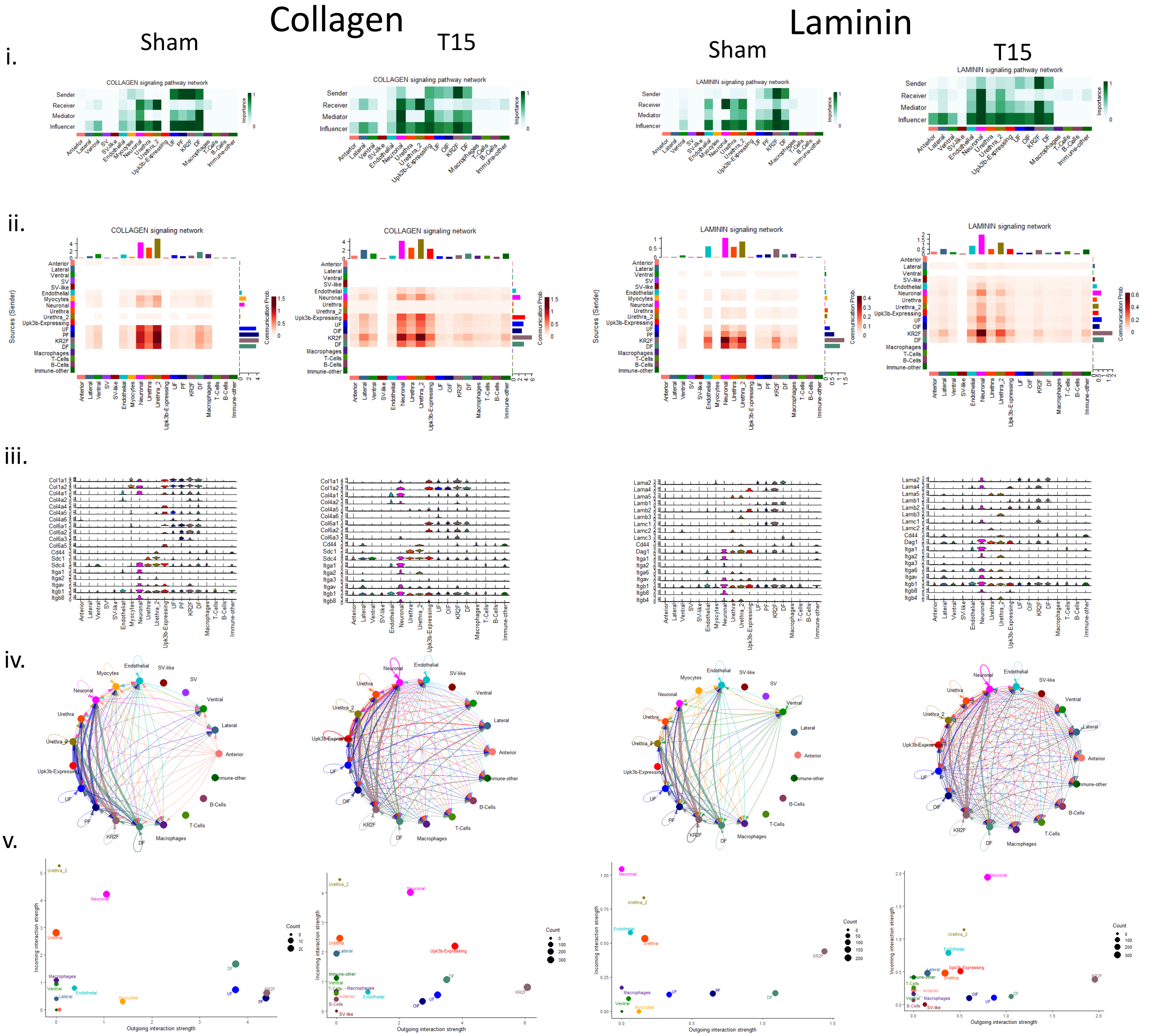

### Supplemental Figure14C

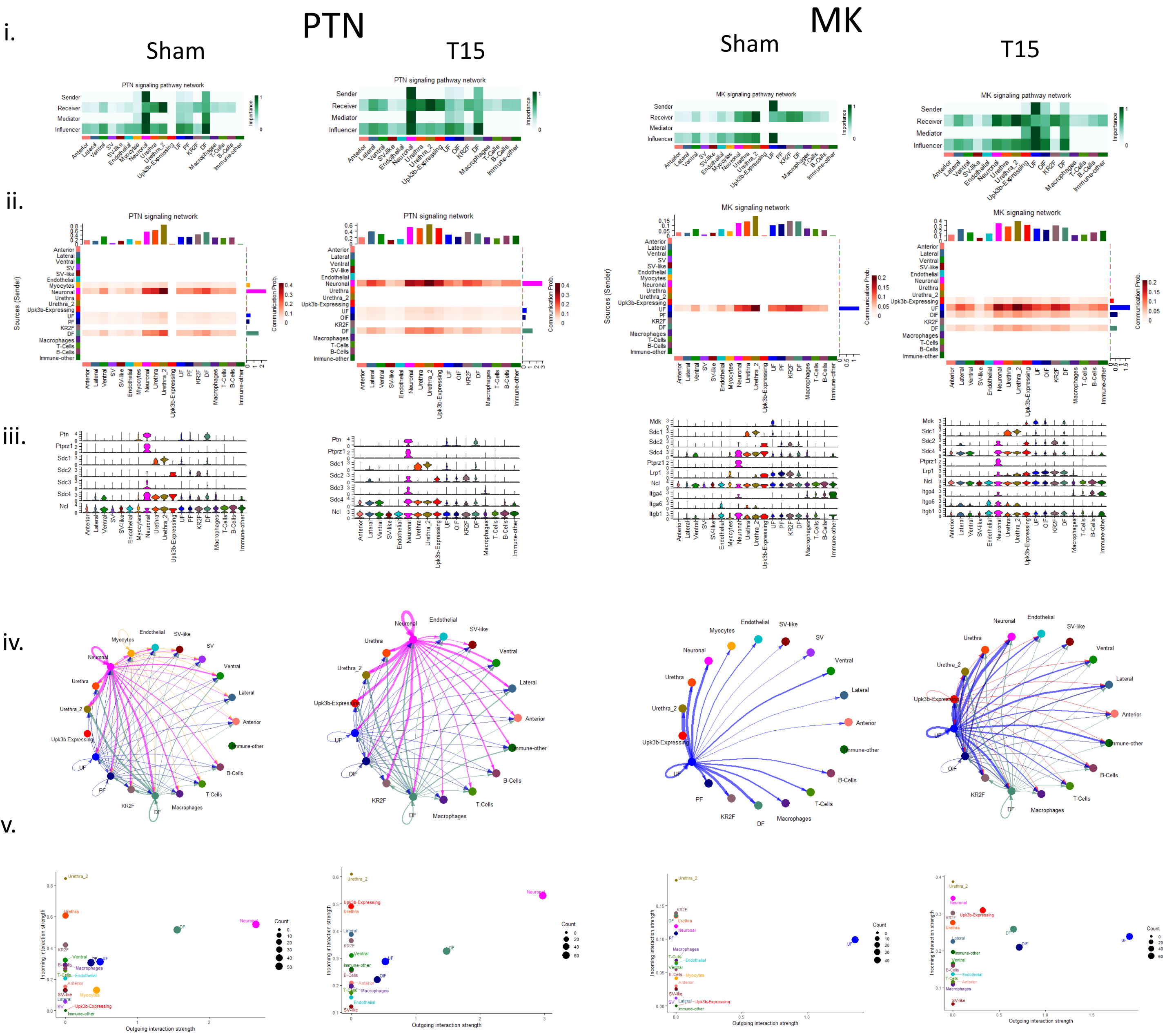

### Supplemental Figure 14D

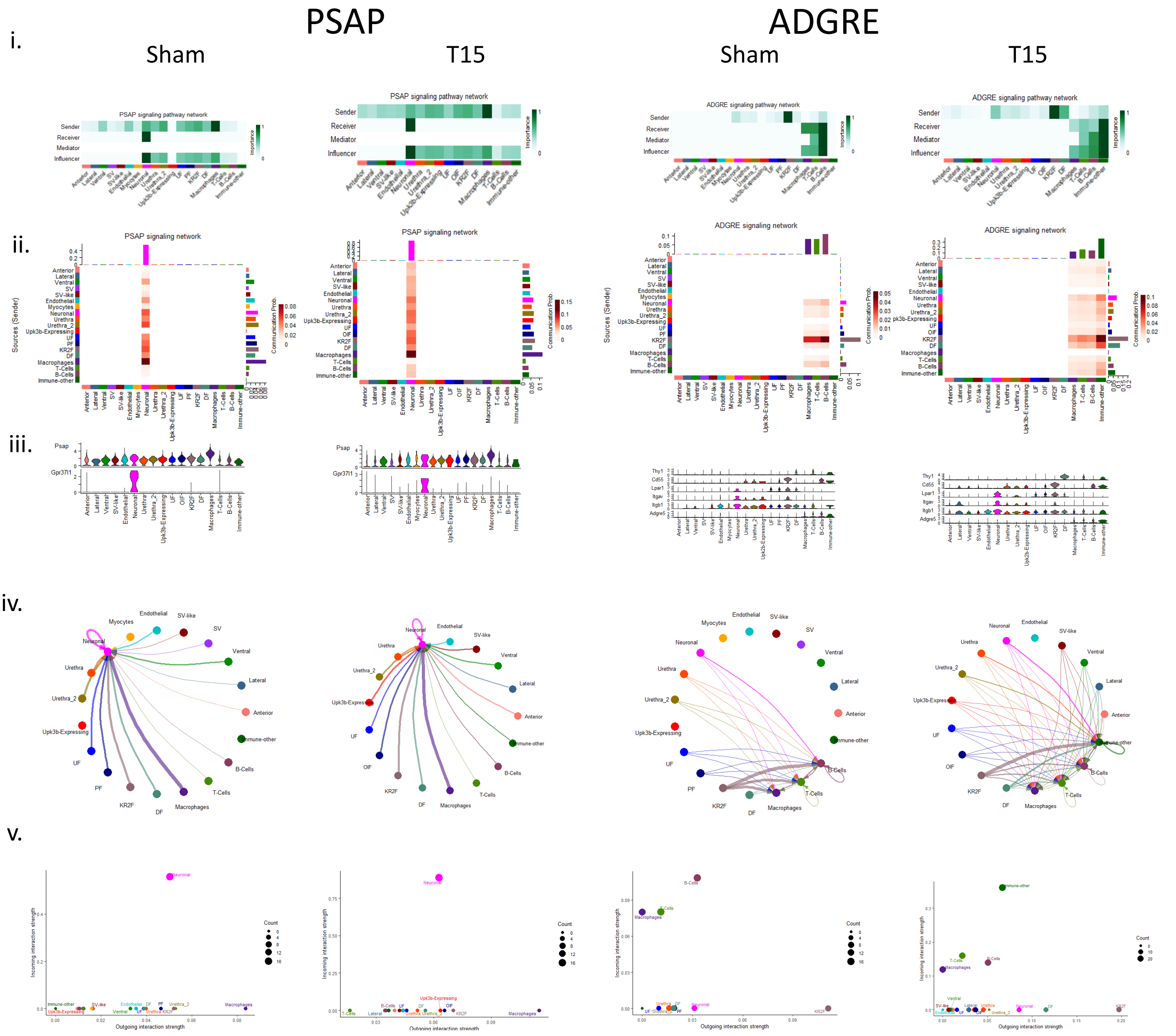

### Supplemental Figure 14E

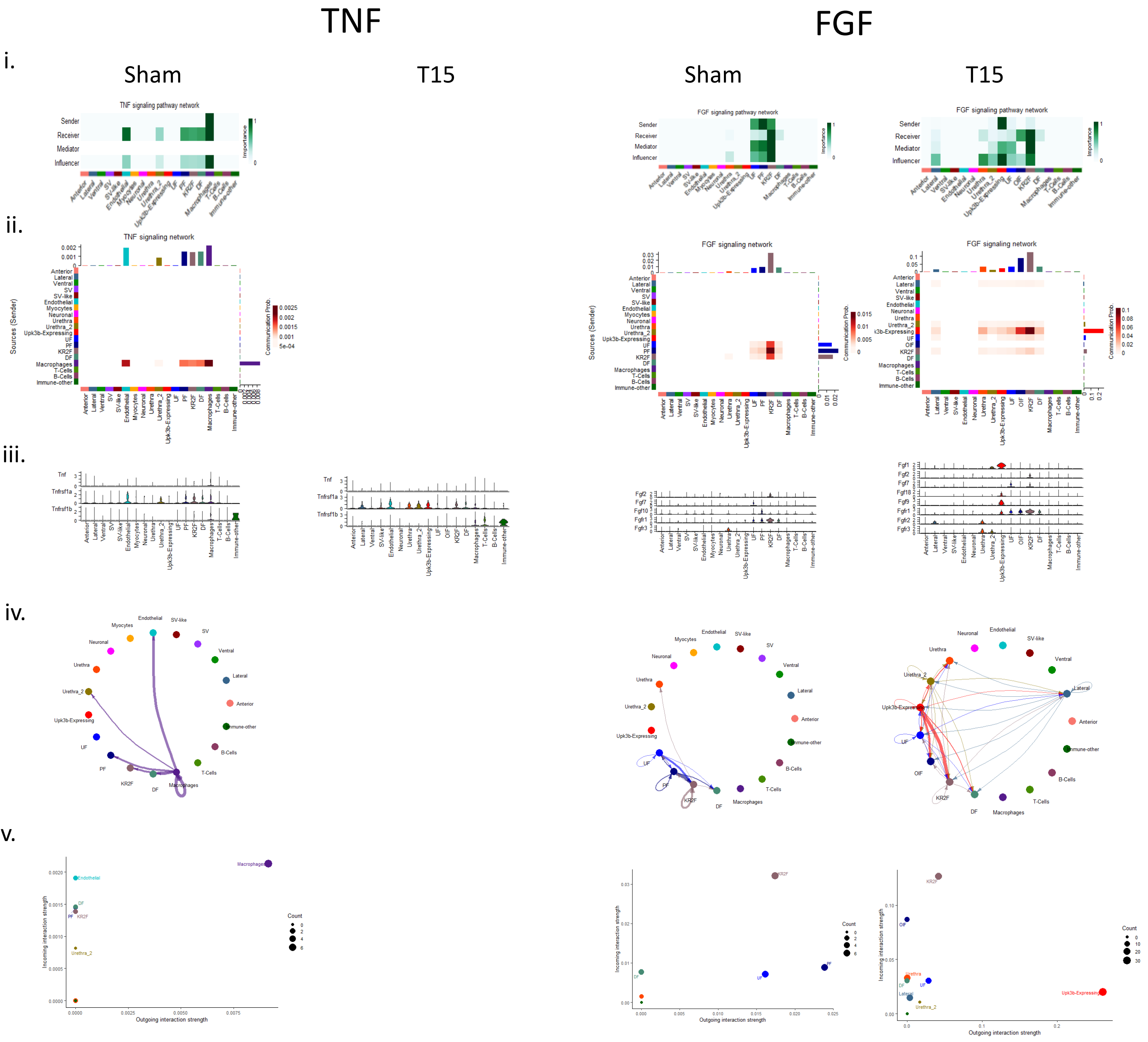
